# Impact of serotonin transporter deficiency on parvalbumin- and neuropeptide Y-producing interneurons of the basolateral amygdala

**DOI:** 10.1101/2025.10.07.680627

**Authors:** Henning Schwert, Elif Salur, Mirjam Richard, Mara Pöllmann, Klaus-Peter Lesch, Esther Asan, Angelika Schmitt-Böhrer

**Affiliations:** Institute of Anatomy and Cell Biology, University of Wuerzburg, Germany; Department of Psychiatry, Psychosomatics and Psychotherapy, Center of Mental Health, University Hospital of Wuerzburg, Germany; Division of Molecular Psychiatry, Center of Mental Health, University of Wuerzburg, Germany; Department of Child and Adolescent Psychiatry, Psychosomatics and Psychotherapy, Center of Mental Health, University Hospital of Wuerzburg, Wuerzburg, Germany; Department of Psychiatry and Neuropsychology, School of Mental Health and Neuroscience, Maastricht University, Maastricht, The Netherlands

**Keywords:** Basolateral amygdala, inhibitory circuits, monoaminergic systems, anxiety, 5-HTT knockout mice

## Abstract

Hyperactivity of the basolateral amygdaloid nuclear complex (BLA) is a hallmark of anxiety-related disorders in humans. Excitation of BLA projection neurons (PN) is fine-tuned by inhibitory interneurons (INs). Monoaminergic afferents to the BLA modulate PN and IN activity. In the present study, BLA-INs immunoreactive(ir) for parvalbumin (PV) or neuropeptide Y (NPY) and their interrelations with serotonergic and catecholaminergic afferents were analyzed in wildtype (WT) and serotonin transporter knockout (5-HTT KO) mice, a model for anxiety- and stress-related disorders. In WT mice, PV- and NPY-ir INs fall into morphological subgroups which possess perisomatic appositions by serotonergic and tyrosine hydroxylase-ir afferents. Dual immunolabeling shows no colocalization of PV and NPY. NPY/somatostatin(SOM) dual labeling documents colocalization of the peptides in some neurons, and single labeling for NPY or SOM in others. These features appear largely preserved in 5-HTT KO mice. However, quantification of PV- and NPY-ir neurons documents a reduction in number and density of NPY-ir neurons throughout the rostrocaudal extent of the amygdala in 5-HTT KO mice. PV-ir neurons remain unchanged. Quantitative PCR shows increased expression of Npy receptor 2, Som receptor 4, and corticotropin releasing factor receptor 1 in the BLA of 5-HTT KO mice. mRNA for the three peptides is unchanged, indicating that it may be NPY propeptide translation which is reduced in 5-HTT KO mice. Taken together, the results document an effect of life-long serotonin imbalance on the BLA NPY-system, which may contribute to previously observed morphological alterations in BLA PNs and increased anxiety-like behavior in 5-HTT KO mice.

## Introduction

The amygdaloid complex plays a central role in the telencephalic processing of emotionally relevant, especially anxiety and fear-eliciting stimuli (LeDoux 2003; Pape et al. 2010; Tovote et al. 2015). Amygdalar hyperactivity and responsitivity are hallmarks of affective disorders (Drevets et al. 2003; Brunetti et al. 2010; Brooks and Stein 2015; Shakman et al. 2016). Hyperexcitation or hyperexcitability of principal neurons (PN) in the lateral (La) and basolateral (BL) nuclei of the amygdala, commonly referred to as the basolateral amygdala (BLA; McDonald, 2023), are thought to be a key feature of amygdalar hyperactivity (Almeida-Suhett et al. 2014; Tovote et al. 2015; Prager et al. 2016; Hinojosa et al. 2022). The BLA nuclei are derived from the ventral pallium (Puelles et al. 2019) and contain a majority of glutamatergic, pyramidal PNs and a minority of non-pyramidal neurons (NPN; McDonald, 1984; Spampinato et al. 2011). Most NPNs are GABAergic interneurons (INs) which fine-tune the activity of PNs (Spampinato et al. 2011; Krabbe et al. 2018; Hajos 2021). INs of the BLA (approximately 15 – 20% of all neurons in the La, 20-25% of all neurons in the BL in rodents; Sah et al. 2003; Hajos 2021; Vereczki et al. 2021) comprise various subpopulations characterized by the separate, sometimes overlapping, expression of the calcium-binding proteins parvalbumin (PV), calbindin (CB), and calretinin (CR), and of neuropeptides such as cholecystokinin, vasoactive intestinal polypeptide, somatostatin (SOM) and neuropeptide Y (NPY) (Asan 1998; McDonald 1989; McDonald and Pearson 1989; Müller et al. 2007a; Pitkänen and Kemppainen 2002; Rainnie et al. 2006; Vereszki et al. 2016, 2021; Hajos 2021). PV-INs are GABAergic and comprise basket cells, whose axons form synaptic contacts on proximal dendrites and somata of PNs, and axo-axonic cells (chandelier cells) which innervate PN axon initial segments (McDonald and Betette, 2001; Müller et al. 2005; Müller et al. 2006; Bienvenue et al. 2012; Krabbe et al. 2018; Vereczki et al. 2016, 2021; Hajos 2021). Optogenetic studies showed an important role of BLA-PV-INs for fear learning in mice (Wolff et al. 2014); a loss of BLA-PV-INs was found to be sufficient to alter behavioral states, such as increasing avoidance behavior and impairing fear learning (Colmers et al. 2025). NPY-producing neurons comprise at least three subpopulations: Firstly, GABAergic INs which most often coexpress SOM and display electrophysiological similarities with SOM-producing INs; secondly, a small group of GABAergic projection neurons also possibly coexpressing SOM (McDonald 1989; McDonald and Pearson 1989; McDonald et al. 2012), and thirdly GABAergic neurogliaform cells which do not produce SOM (Vereczki et al. 2021; Ozsvar et al. 2024). In the rat BLA, NPY- and PV-immunoreactivity (IR) is mutually exclusive (McDonald 2023). In contrast, a recent study in mice using transgenic and viral strategies suggested that about a fourth and a third of NPY-producing BLA neurons are PV- or SOM-expressing INs, respectively, and that almost half are neurogliaform cells which do not produce SOM or PV (Vereszki et al. 2021). SOM-INs, presumably including those that additionally produce NPY, are Martinotti cells forming inhibitory synapses on distal dendritic segments of PNs. In contrast, neurogliaform cells are thought to release GABA predominantly via volume transmission, i.e. from axonal varicosities without a synaptic partner (Ozsvar et al. 2021). Released GABA activates postsynaptic and extrasynaptic GABA_A_- and GABA_B_-receptors on PNs, triggering a biphasic mixture of fast and slow inhibition (Ozsvar et al. 2024). In rodents, suggested effects of NPY in the BLA *in vivo* include the induction of short- and long-term anxiolysis, and NPY in the BLA is thought to be crucially involved in the regulation of stress- and anxiety-related behavior (Kask et al. 2002; Yilmazer-Hanke et al. 2002, 2004; Gutman et al. 2008; Truitt et al. 2009; Cohen et al. 2012; Tasan et al. 2016; Bompolaki et al. 2024). In humans, a 30% decrease in NPY mRNA expression, due to a genetic polymorphism within the NPY promoter region, is associated with increased amygdala activity upon exposure to threat-related facial expressions (Zhou et al. 2008). Anxiolytic effects of NPY in the BLA appear to be mediated by activation of NPY receptor 1 (NPYR1) which *in vitro* has been shown to produce a direct hyperpolarization-induced inhibition of PNs. This can be countervailed by corticotropin releasing factor (CRF), a potently anxiogenic peptide (Giesbrecht et al. 2010). Somewhat counterintuitively, selective activation of the mainly presynaptic NPYR2 in the BLA has acutely anxiogenic effects (Nakajima et al. 1998; Sajdyk et al. 2002).

Information processing in the BLA is modulated by monoaminergic afferents (Asan 1998; Asan et al. 2013; Bocchio et al. 2016; McDonald 2023). Dopaminergic and serotonergic innervation from the substantia nigra/ventral tegmental area (SN/VTA) and the dorsal raphe nucleus (DRN), respectively, are thought to exert differential effects on BLA neuronal circuits, enabling flexible behavioral responses to variable environmental requirements, for instance in the course of the stress response (Asan et al. 2013; Kroner et al. 2005; Bocchio et al. 2016). Dysregulation of monoaminergic influence on BLA activity is thought to contribute to stress-related disorders in patients, and therapeutic interventions directed at monoaminergic transmission modulate amygdala activity as shown by fMRI studies (Godlewska et al. 2016). Therefore, neural processes underlying the actions of these transmitters in the BLA have been the subject of numerous investigations (reviewed in McDonald 2023). Due to the crucial role of INs for fine-tuning BLA-PN activity, the monoaminergic modulation of inhibitory circuits is of particular interest (Bissiere et al. 2003; Bocchio et al. 2015; Prager et al. 2016). In rats, synaptic contacts of dopaminergic and serotonergic axonal varicosities both on PNs and on INs in the rat BLA have been documented (Müller et al. 2007b, 2009; Bonn et al. 2013). Additionally, monoaminergic afferents are thought to also signal via volume transmission (Gianni and Pasqualetti, 2023; McDonald 2023). Released transmitter molecules diffuse in the extracellular space for a variable distance, depending on the texture of the extracellular matrix and on the presence of transmitter metabolizing enzymes or membrane transporters which remove the transmitters from the extracellular compartment. Effects in target cells will depend on the concentration of transmitter molecules at a given distance from the exocytosing varicosity, and on the affinity of the receptors expressed by the target cell. Maximal effects are to be expected if varicosities and target membranes are in close apposition. Electron microscopic studies have documented both post- and extrasynaptic localization of dopamine receptors in the BLA (Pickel et al. 2006) and extrasynaptic localization of serotonin (5-HT) receptors in cortical areas (Riad et al. 2000), indicating that both synaptic and volume transmission mediate effects of these monoamines in the telencephalon (McDonald 2023). Furthermore, catecholamines, particularly dopamine, and the indolamine 5-HT play important roles in brain development, and considerable evidence links dysfunction of monoaminergic systems to neurodevelopmental disorders (Money and Stanwood, 2013; Lesch and Waider, 2012). The serotonergic system is of particular clinical relevance, since numerous genetic variants in 5-HT transmission molecules have been identified which may influence individual differences in the stress response and in the risk for stress-related disorders (Holmes 2008). In humans and non-human primates, a functional polymorphism in the transcriptional control region of the 5-HT transporter (*5-HTT, SLC6A4*) gene, which leads to reduced production of 5-HTT in carriers of the short allele, has been shown to interact with early-life adversity to convey an increased risk for affective dysregulation (Lesch et al., 1996; Holmes, 2008; Asan et al. 2013; Homberg et al. 2016). The 5-HTT knockout (KO) mouse, generated to investigate the neural basis of these phenomena (Bengel et al. 1998), displays an increased anxiety-like behavior and has become a widely accepted mouse model for investigations into the neurobiological basis of anxiety- and stress-associated diseases (Holmes et al. 2003a,b; Kim et al. 2005; Holmes 2008). BLA-PN in these mice display increased dendritic spine density compared to wildtype (WT) mice, presumably representing a morphological correlate of hyperexcitability, and the animals display deficits in fear extinction recall (Wellmann et al. 2007; Nietzer et al. 2011). In 5-HTT overexpressing mice, which are characterized by a low anxiety phenotype (Jennings et al. 2006) and impaired fear memory (Barkus et al. 2014), the recruitment of PV-INs is impaired during fear, a phenomenon mediated by reduced functionality of the 5-HT2A receptor expressed on these neurons (Bocchio et al. 2015). Alterations in BLA inhibitory circuits have not been studied in 5-HTT KO mice to date. In view of the finding that SN/VTA dopaminergic neurons which provide innervation of the BLA are densely innervated by serotonergic DRN neurons, it is also of interest whether the 5-HTT deficiency influences the catecholaminergic, particularly the dopaminergic innervation of the BLA. Therefore, the present study was designed to investigate the morphology, distribution and density of PV- and NPY-INs in the BLA, to assess their interrelations with monoaminergic afferents, and to study the effects of life-long 5-HT imbalance in 5-HTT KO mice on these parameters. Additionally, mRNA expression levels of *Npy* and its receptors *Npyr1* and *Npyr2* in the BLA of WT and KO mice were studied, as were the mRNA expression levels of other molecules involved in mediating anxiolytic or anxiogenic effects in the BLA, namely *Som* and its receptors (*Sstr2, Sstr4*), *Crf* and its receptors (*Crfr1, Crfr2*), and 5-HT receptors (*5Htr1a*, *5Htr2a*, and *5Htr2c*) (Asan et al. 2013; Giesbrecht et al. 2010; Scheich et al. 2016).

## Materials and methods

### Animals

Experiments were carried out on adult (>10 weeks), male mice of a 5-HTT KO breeding line backcrossed into a C57BL/6J background for >10 generations (Bengel et al. 1998). The animals were either homozygous for the wild-type (WT) allele or homozygous for the KO allele. Water and food were available *ad libitum*. All experiments were carried out according to the European Parliament and Council Directives (86/609/EEC and 2010/63/EU) and in accordance with the Guide for the Care and Use of Laboratory Animals of the Government of Lower Franconia (approval code 55.2-2531.01-93/12). The number of animals and the suffering induced was always kept as low as possible.

### Tissue preparation

For the single and dual immunolabeling studies mice were sacrificed by anaesthetic overdosing and transcardially perfused using 4% paraformaldehyde in 0.01 M phosphate buffered saline pH 7.4 (PBS). Brains were dissected, postfixed in the same fixative overnight, washed in PBS, cryoprotected in 10% and 20% sucrose in PBS, frozen in liquid nitrogen-cooled isopentane and stored at −80°. For sectioning, brains were either slowly thawed to room temperature (RT), coronal 35 µm-thick sections through the entire rostro-caudal extent of the amygdaloid complex were cut in PBS on a Vibratome (Leica, Wetzlar, Germany), collected in five series and used either immediately for immunolabeling or stored in cryoprotectant solution (Asan 1998) at −20°C. One series of sections was mounted on Superfrost^TM^ microscopic slides, and subjected to Gallocyanin staining (Heinsen et al. 2000). Alternatively, frozen brains were cut at 18 µm thickness on a cryostat (Leica, Wetzlar, Germany), thawed onto HistoBond® adhesive microscope slides (Marienfeld) additionally pre-coated with glycerin. Slides were stored at −80°C until being used for immunolabeling.

For the gene expression study animals were sacrificed by cervical dislocation under deep anesthesia with isoflurane. Brains were then dissected and frozen in dry ice-cooled isopentane and stored at −80°C until further processing.

### Immunolabeling

#### For light microscopic (LM) immunoenzyme single labeling

After thorough washing in 0.01M phosphate-buffered saline pH 7.4 (PBS), free-floating vibratome sections were preincubated for two hours at room temperature in PBS with 1% Triton X-100 and 5% normal goat serum (NGS; blocking solution). Two adjacent series of vibratome sections were incubated in monoclonal mouse anti-PV IgG (Millipore, #MAB1572; diluted 1:4000) and polyclonal rabbit anti-NPY IgG (Sigma-Aldrich, #N9528; diluted 1:8000) in PBS with 0.5% Triton X-100 (Sigma), 1% NGS (incubation solution). For primary antibody incubation, 0.05%NaN_3_ was added to the incubation solution. Monoaminergic innervation was assessed in further section series using anti-tyrosine hydroxylase (TH; rabbit, polyclonal, Millipore, #AB152, 1:500), anti-5-HT (rabbit, Sigma, #S5545, or Immunostar, #20080, 1:2000 or 1:15000-1:40000), and anti-5-HTT (rabbit, polyclonal, Calbiochem, #PC177L, 1:500). Sections were incubated in primary antibodies for 48-72 hours at 4°C, washed in PBS, incubated overnight at 4°C in biotinylated goat anti-mouse IgG or biotinylated goat anti-rabbit IgG secondary antibodies in incubation solution as required (Vector Laboratories, mouse: #BA-9200, 1:500; rabbit: #BA-1000, 1:500), washed again in PBS, incubated in avidin-biotin–horseradish-peroxidase complex (DAKO) prepared as decribed by the manufacturer in PBS for 2 hours, and washed again in PBS. Visualization was carried out using the glucose-oxidase-diaminobenzidine (DAB) method described by Zaborszky and Heimer (1989). Immunolabeling on the majority of sections was intensified using nickel intensification (NiDAB; Asan 1997). Immunoreaction product development was checked by LM and stopped by repeated washes in PBS when maximum specific labeling was achieved. Sections were either washed in PBS and processed for dual immunolabeling as described below, or mounted onto Superfrost^TM^ (Menzel, Braunschweig, Germany) microscopic slides, dried, dehydrated in a graded ethanol series and mounted in DePex (Serva, Heidelberg, Germany) from xylene. For *dual immunoenzyme labeling*, an established sequential incubation protocol was used (Eliava et al., 2003): Anti-TH, anti-5-HTT or anti-5-HT reacted sections developed using Ni-DAB were subjected to a second immunoreaction for detection of NPY or PV as described above, omitting the preincubation step. Immunolabeling was visualized using non-intensified DAB (Eliava et al. 2003).

*Single and dual fluorescence immunolabelings* were carried out by incubation of free-floating vibratome sections as described above with one primary antibody, or by simultaneous incubation of sections with two primary antibodies generated in different species, namely rabbit anti-5-HTT, mouse anti-PV, rabbit anti-NPY and rat anti-SOM (monoclonal, clone YC7, Sigma-Aldrich, #MAB354, 1:500). After washing in PBS, the sections were incubated with the appropriate fluorochrome-labeled secondary antibody or secondary antibody combination (Cy2- or Alexa488-labeled anti-mouse, anti-rat IgG or anti-rabbit IgG and Cy3- or Alexa555-labeled anti-rat or anti-rabbit IgG; Vector Laboratories or Invitrogen; dilution 1:200, 1:400 or 1:600). Secondary antibody incubations and subsequent washing in PBS was carried out as described above while keeping the sections in the dark. Mounted cryosections were washed in PBS, incubated in blocking solution for 2 hours, incubated with anti-NPY and/or anti-SOM (antibody dilution 1:500) in incubation solution (with 0.3% Triton-X-100) overnight at 4°C in a humid chamber, washed again in PBS and incubated in secondary antibody solution for 2h at room temperature (RT) and washed in PBS. In some sections, an antibody retrieval step (incubation in 10mM sodium citrate with 0.05% TWEEN 20, pH 6.0, for 20 min at 80°C) was performed before preincubation with blocking solution. Fluorescent sections were counterstained using 4’,6-diamidino-2-phenylindole (DAPI). After washing in PBS, the free-floating sections were mounted on Superfrost microscopic slides. All fluorescent sections were covered with Fluoro-Gel (Electron-microscopic sciences, Hatfield, USA) and observed using a Zeiss Axioskop equipped with appropriate filter systems or the Compact Fluorescence Microscope BZ-X800 (Keyence, Neu-Isenburg, Germany).

*Controls* were performed for all single and dual immunolabeling experiments by omitting the primary antibodies for single labeling, by omitting either of the primary antibodies for dual fluorescence labeling, and by omitting the first and/or second primary antibody in sequential immunoenzyme labelings. Incubations with the respective secondary antibodies and, if applicable, Ni-DAB/DAB developments were performed according to the respective protocols. Additionally, individual sections of 5-HTT KO animals were incubated with the anti-5-HTT antibody. None of these preparations showed specific immunolabeling.

### Analysis of appositions between monoaminergic afferents and PV-ir and NPY-ir neurons

Perisomatic appositions were analyzed according to previously established methods (Bonn et al. 2013). Dually DAB-immunolabeled amygdala sections were observed using a 100x oil objective. Immunostained neurons containing a nucleus were randomly selected and categorized morphologically (see below). The somata of labeled neurons were studied throughout all focal levels. Appositions of 5-HTT, 5-HT and TH-immunoreactive (-ir) fibers on PV or NPY-ir somata, in which no gap between the differently labeled structures was discernible, were regarded as appositions. For quantitative evaluations, perisomatic appositions were counted in dually labeled coronal sections at different interaural levels (−0.80 mm to −2.9 mm from Bregma; Franklin and Paxinos, 1997) from eight to nine WT (5-HTT/PV-, 5-HTT/NPY-, TH/PV- and TH/NPY-labeled sections) and from eight 5-HTT KO mice (TH/PV- and TH/NPY-labeled sections). Appositions were recorded separately for different morphological types of PV- and NPY-ir neurons in the different nuclei and subnuclei of the BLA. For type I PV-ir neurons in the BL (see below), mean apposition counts (from n≥3 neurons in each of 6 animals per group) were determined for 5-HTT-(in WT) and TH-ir appositions (in WT and 5-HTT KO mice). Statistically significant differences were absent between the individual animals of each group in this category, indicating similar labeling quality of dopaminergic or serotonergic afferents in the sections analyzed. Apposition counts were therefore pooled for mice of the respective genotypes in each category. Only categories with ≥4 neurons from ≥3 animals were used for statistical comparisons.

### Morphometric analyses of immunoenzyme reacted sections

The number of PV- or NPY-ir neurons in La and BL was analyzed in six coronal sections from eight to nine WT and eight 5-HTT KO mice by an observer blinded for the respective genotype. The sections were spaced 140 µm apart and covered the rostro-caudal extent of the amygdala, with the two rostral-most sections (−0.80 to −1.5 mm from bregma) representing anterior amygdaloid levels, two middle sections (−1.5 to −2.2 mm) representing intermediate, and two caudal sections (−2.2 to −2.9 mm) representing posterior levels of the amygdala. All immunoreactive neurons of the La and BL displaying a nucleus were counted in each section. La and BL boundaries and, additionally, in the BL the anterior and posterior subdivisions (BLa, BLp) were delineated in each section according to the “Mouse Brain in Stereotaxic Coordinates” (Franklin and Paxinos, 1997). Delineation in difficult cases was supported by comparison with Gallocyanin-stained adjacent sections, and immunolabeled neuron numbers, positions within the La/BL and borders of the respective nuclei were recorded using the “Stereo Investigator” software (version 10.52) and the Neurolucida explorer (version 10.52; MicroBrightfield Europe B.V., the Netherlands). Neuron density in the different nuclei, subnuclei and rostro-caudal regions was calculated by dividing the number of neurons counted by the respective amygdalar nuclear volume analyzed (area analyzed x section thickness) and depicted as number/0.001mm^3^.

### Gene expression analyses

Gene expression analyses were carried out on ten male 5-HTT WT and eight male 5-HTT KO mice. For Laser capture microdissection, Brains were sectioned at 20 µm in a Leica CM1950 freezing microtome at −20°C and the tissue sections were mounted on polyethylene naphthalate (PEN) membrane slides 2.0 µm (Leica, Wetzlar, Germany). Mounted tissue sections were fixed in −20°C cold 75% EtOH (diluted in DEPC-treated water) for 2 min, stained with 0.25% cresyl violet and then processed through an ascending ethanol series (75%, 90%, 100% ethanol for 20 s each) for decoloring. After an additional incubation for 1 min in 100% ethanol, the slides were air dried in 50 ml tubes with Silica Gel Orange (Roth, Karlsruhe, Germany), and the tissue was dissected using a Leica LMD6 laser microdissection system, which allowed the precise extraction of the La/BL (−0,94 to −1,82 from bregma).

RNA was extracted from the dissected tissue using the miRNeasy Micro Kit (Qiagen, Hilden, Germany) and stored at −80°C until further use. The iScript cDNA Synthesis Kit from Bio-Rad was utilized for Reverse transcription of the RNA into cDNA. As the LMD only results in low amounts of harvested tissue, extracted RNA and subsequent cDNA, an unbiased, target-specific preamplification of cDNA using the 2x SsoAdvanced PreAmp Supermix (Bio-Rad) was necessary. Primer sequences to detect the expression levels of genes related to NPY, SOM, CRF, and the serotonergic system required for this pre-amplification step as well as for the quantitative real-time PCR (qPCR), were selected using the NCBI BLAST tool (Online resource 1) and purchased from Sigma-Aldrich (St. Louis, MO, USA). To perform the qPCR reactions, the pre-amplified cDNA stock solution was further diluted using water to a 1:20 working solution. Per sample, 0.5 µl forward and reverse primer (5 µM each) were mixed with 5µl SYBR™ Select Master Mix (Thermo Fisher Scientific, Waltham, MA, USA) and 4 µl cDNA to a total volume of 10 µl. A 384-well plate in a CFX384 real-time PCR detection system (Bio-Rad) was used for the reactions. LinRegPCR program (version 2017) (Ruijter et al., 2009) was used to calculate mean PCR efficiencies. Evaluation of expression, normalization and calculation of relative gene expression results were obtained with qBase+ software (Biogazelle, Ghent, Belgium)

### Statistics

For statistical evaluation of the immunolabeling data and creation of respective graphs the software GraphPad Prism version 5.03 (GraphPad Software, La Jolla, California, USA) was used. Group comparisons were performed using the non-parametric Mann-Whitney-U test. Gene expression data sets were analyzed using SPSS Statistics version 26 (IBM Corporation, New York, USA) and GraphPad Prism software version 10 was used for data graphing. A Levene test for homogeneity of variances and a Shapiro-Wilks test for normal distribution were performed on these data. If distributions of variance were not significant (p > 0.05) and samples where normally distributed a t-test was performed. In case the requirements for a t-test for independent samples were not met, the Mann-Whitney U-test was used for pairwise comparisons. Statistical significance was set at p < 0.05. Significance levels are indicated (if not otherwise indicated in the graph legends) as * p < 0.05, ** p < 0.01, *** p < 0.001, **** p < 0.0001.

## Results

### Distribution and morphology of PV- and NT-ir neurons in WT and 5-HTT KO mouse BLA

#### PV-ir neurons

In both WT and 5-HTT KO mice, immunoenzyme labeling detects the highest density of PV-ir neuronal somata in the BLp, lower density in the BLa, lowest in the La (Fig. 1a,b). Also in both genotypes, PV-ir neurons in the BLA fall into two groups based on soma shape and dendritic morphology: the larger group are multipolar neurons with three to five dendrites and round to oval somata of varying size. In analogy to the morphological types of BLA PV-neurons in the rat (Kemppainen et al. 2000), this group can additionally be divided into two subgroups, type I with comparatively small and type II with comparatively large somata. A further group, type III, consists of neurons with more fusiform somata and dendrites originating mainly from the two cell poles (Fig. 1c,d). The reaction product is homogeneously distributed in somata, proximal dendrites and axon hillocks. The intermediate and distal dendrites are not commonly labeled. The proximal dendrites of type I and type II neurons display mostly smooth contours. The proximal dendrites of type III neurons are also smooth but often narrower than those of type I and -II neurons.

**Fig. 1.**
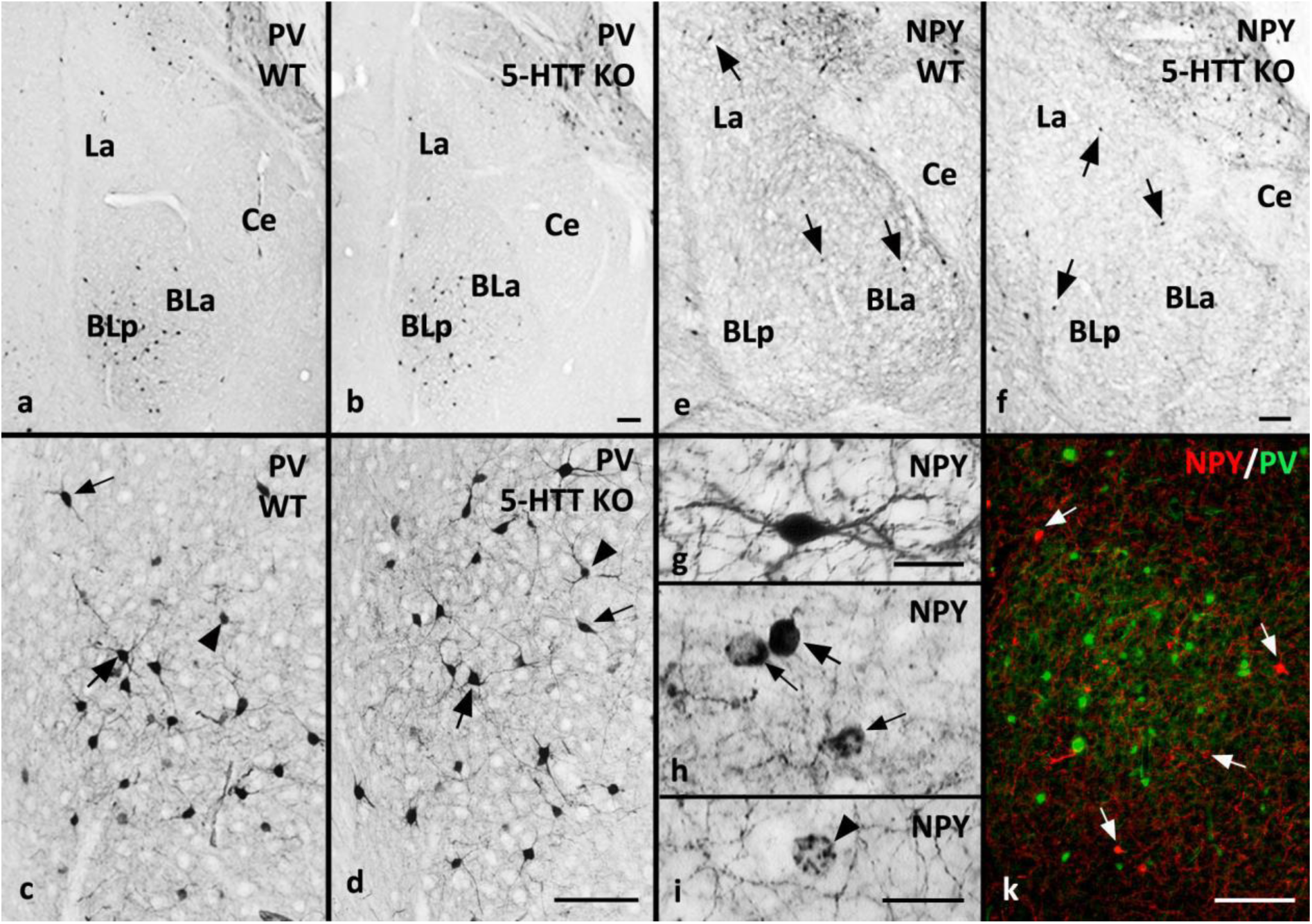
Immunoenzyme labeling for PV in the WT **(a)** and 5-HTT KO mouse BLA **(b)**. In mice of both genotypes, PV-ir neurons are densely clustered in the posterior basolateral nucleus (BLp), less dense in the anterior BL (BLa), and least dense in the lateral nucleus (La). **(c,d)** Type 1 (large arrows), II (arrowheads) and III (small arrows) PV-ir neurons in the WT (**c**) and 5-HTT KO mouse (**d**). **(e,f)** Immunoenzyme labeling for NPY in the WT **(e)** and 5-HTT KO mouse **(f)**. Neurons are scarce but evenly distributed in the BLA of both genotypes. **(g)** Paradigm fusiform and **(h)** round NPY-ir neurons. **(h)** NPY-ir neurons show different labeling intensities, occasionally NPY immunoreaction product labels the perikaryon homogeneously (large arrow), in other cases NPY-IR is localized in “cytoplasmic streets” (small arrows). **(i)** Occasionally, NPY-ir elements appear to surround an unlabeled neuron. **(k)** Dual immunofluorescence labeling for NPY (red, some neurons designated by arrows) and PV (green) show non-overlapping immunoreactivity for the two markers in the BLA. Ce: central amygdaloid nucleus. Bars: 100 µm in b for a, b, in d for c, d, in f for e, f, and in k; 20 µm in g for g, and in i for h, i.

#### NPY-ir neurons

In both genotypes, NPY-ir neurons are less frequent and appear more homogeneously distributed throughout the La and BL than PV-ir neurons (Fig. 1 e,f). BLA-NPY-ir neurons can be divided into two classes based on soma shape: neurons with round and neurons with fusiform somata (Fig. 1g,h). The immunoreaction product is homogeneously distributed throughout the somatic cytoplasm in many neurons (Fig. 1 g,h), but forms “cytoplasmic streets” in others (Fig. 1h); labeling only occasionally extends into proximal dendrites and axons (Fig. 1g). Immunolabeled proximal dendrites are narrow and smooth. Most NPY-ir fiber plexus are constituted by narrow fibers with numerous small varicosities. Occasionally, NPY-ir varicose structures were observed surrounding somata of apparently non-immunoreactive neurons (Fig. 1i). In these cases, it was often possible to trace the varicose structures to parent axons when viewed through different focus levels.

### Colocalization of PV- or SOM-immunoreactivity in NPY-ir BLA neurons in WT and 5-HTT KO mice

In both genotypes, we did not observe PV/NPY-double-labeled cells in the BLA (Fig 1k). In contrast, NPY/SOM-double-labeled and SOM- or NPY-single-labeled neurons are found in La and BL (Fig. 2, Fig. 3). NPY-ir single-labeled cells are often small and round, SOM-ir single labeled and SOM/NPY double labeled cells appear larger and often more ovally shaped (Fig. 2). The relative immunofluorescence intensity for the two markers differs between double-labeled neurons. In some double-labeled neurons, immunolabeling for NPY and SOM is homogeneous throughout neuronal perikarya and extends occasionally also into dendrites (Fig. 3a-i). More often, immunofluorescence is concentrated in the perinuclear cytoplasm. Perikaryal IR for NPY and SOM is localized in specific, largely but not in all cases completely overlapping compartments in double-labeled neurons (Fig.3g-m). In some cases, particularly in cryostat sections, it is difficult to discern perinuclear cytoplasmic labeling from terminal axon baskets as shown in the DAB-stained sections (Fig. 3n-p, cf. Fig. 1 i).

**Fig. 2.**
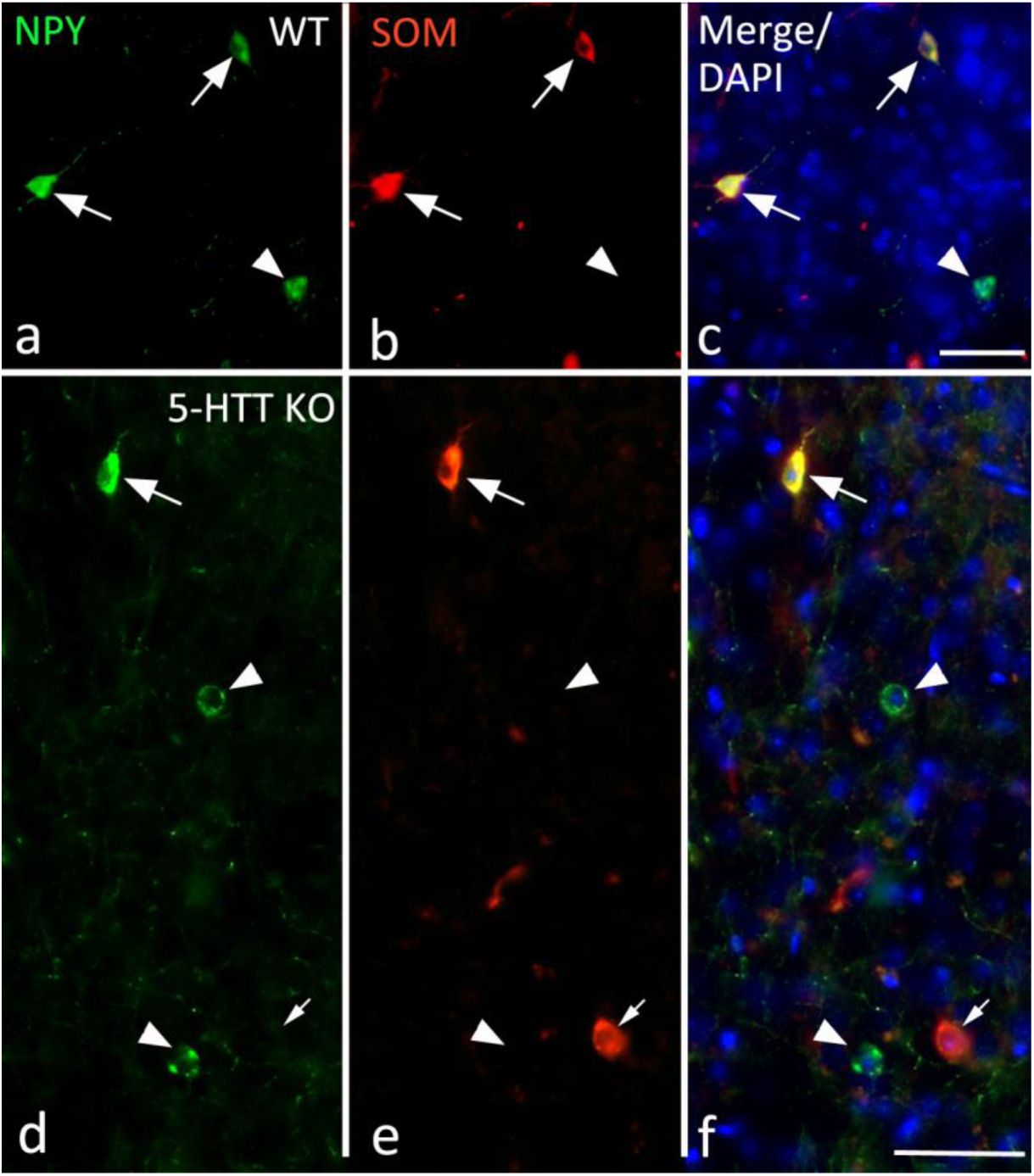
Dual immunofluorescence labeling for NPY and SOM in the BLA of WT **(a-c)** and 5-HTT KO mice **(d-f)** shows double-labeled neurons (arrows) as well as SOM-single labeled neurons (small arrows) and NPY single labeled neurons (arrowheads). Single NPY-ir neurons often possess small round perikarya, double-labeled and single SOM-ir neuron perikaryal are often larger and more ovally shaped. Vibratome sections, Bars = 50 µm.

**Fig. 3.**
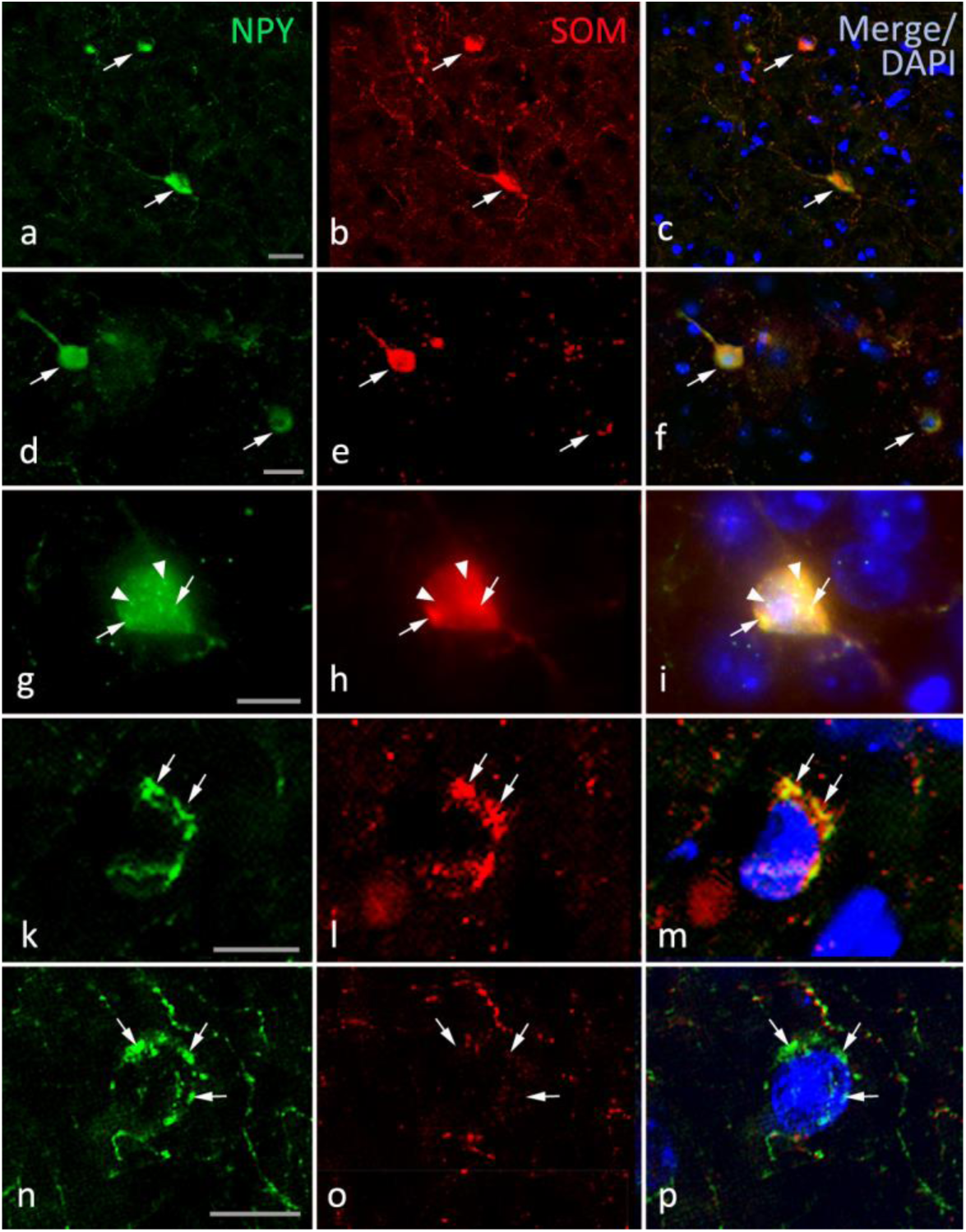
Dua**l** Immunolabeling for NPY und SOM in the BL of WT **(a-c, g-m)** and 5-HTT KO mice **(d-f, n-p)**. **(a-f)** Immunolabeling intensity varies between double-labeled neurons (arrows) both in WT **(a-c)** and in 5-HTT KO mouse BLA **(d-f)**. **(g-i)** SOM and NPY immunofluorescence are homogeneously distributed in the perikaryal cytoplasm and additionally concentrated in overlapping compartments (arrows). Intense NPY-IR appears localized in additional compartments (arrowheads). **(k-m)** SOM- and NPY-IR are localized in overlapping perikaryal compartments (arrows). **(n-p)** Single NPY-IR is found in varicose structures surrounding neuronal nuclei. Differentiation between labeling in perikaryal compartments as opposed to NPY-ir varicose axons surrounding the perikaryon (cf. Fig. 1 h, i) is not always possible. a-f and k-p: cryostat sections, g-f: vibratome section. Bar in a, d: 20 µm, in g, k, n: 10 µm.

### Interrelations between monoaminergic innervation and PV- and NPY-producing BLA neurons in WT and 5-HTT KO mice

#### BLA innervation by 5-HTT and 5-HT fibers

5-HTT immunoreactions in WT result in intra- and interindividually reproducible labeling of dense serotonergic fiber plexus in the BLA (Fig. 4a, Fig. 5a, b). 5-HTT fiber density in 5-HTT/PV (Fig. 4b, 5a) and 5-HTT/NPY dual immunolabeled sections is as high as in single labelings (Fig. 4 a,c). 5-HTT-ir axons display numerous irregular varicosities (Fig. 4b, c). 5-HTT fiber density is higher in the BL than in the La (Fig. 4a) and appears slightly higher in the BLa than in the BLp. In contrast to 5-HTT immunoreactions, 5-HT immunoreactions vary considerably between animals. When 5-HT-ir fiber density is comparable to that of 5-HTT-ir fibers in WT sections (Fig. 4c,d), 5-HT-ir fibers are observed to be fine and tortuous, occasionally with pleomorphic varicosities, but altogether smoother than terminal axons detected in 5-HTT immunoreactions. M-type fibers, characterized by typically large varicosities and low 5-HTT content, are not detected in 5-HT-labeled sections of the La/BL, indicating that 5-HTT immunoreactions quantitatively document the serotonergic afferents to the nuclei. In 5-HTT KO mice, 5-HT immunoreactions also show a high interindividual variability in detection quality. In sections from some 5-HTT KO animals, 5-HT-ir fiber plexus density appears as high as in sections from well-reacted WT animals (cf. Fig. 4, 5). As in WT animals, 5-HT-ir fiber plexus are of higher density in the BL than in the La (not shown).

**Fig. 4.**
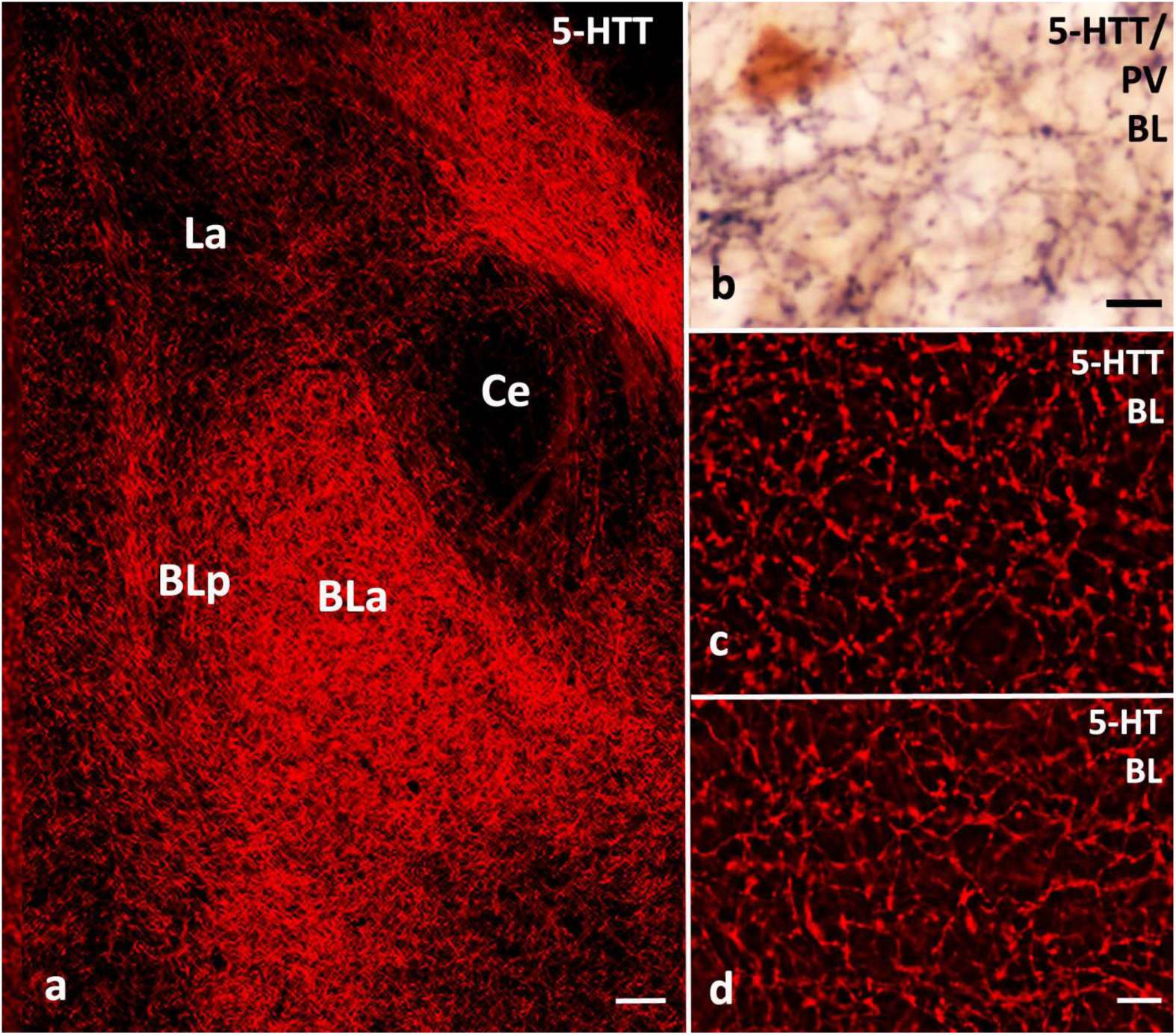
Serotonergic fibers in the WT mouse. **(a)** 5-HTT immunofluorescence shows dense serotonergic fiber plexus in the BLa, somewhat less dense in the BLp and least dense in the the La. **(b)** 5-HTT-ir Fiber density in dual immunoenzyme labeling for 5HTT (black)/PV (brown) is comparable to density in single 5-HTT labelings. 5-HTT-ir fibers show small varicosities forming appositions on a PV-ir neuron. **(c,d)** Immunoenzyme-labeled 5-HTT and 5-HT fibers are shown after digital conversion into false colors. The varicose morphology of 5-HTT fibers (**c**) is more evident. **(d)** Fiber plexus in a 5-HT immunoreacted section with excellent fiber staining. The density of 5-HT-ir fibers is similar to that of 5-HTT-ir fibers. 5-HT-ir fibers contain longer stretches of smoothly contoured fibers than 5-HTT-ir fibers. Bar in (a) represents 100 µm, in (b) 10 µm, and in d for c and d 10 µm.

**Fig. 5.**
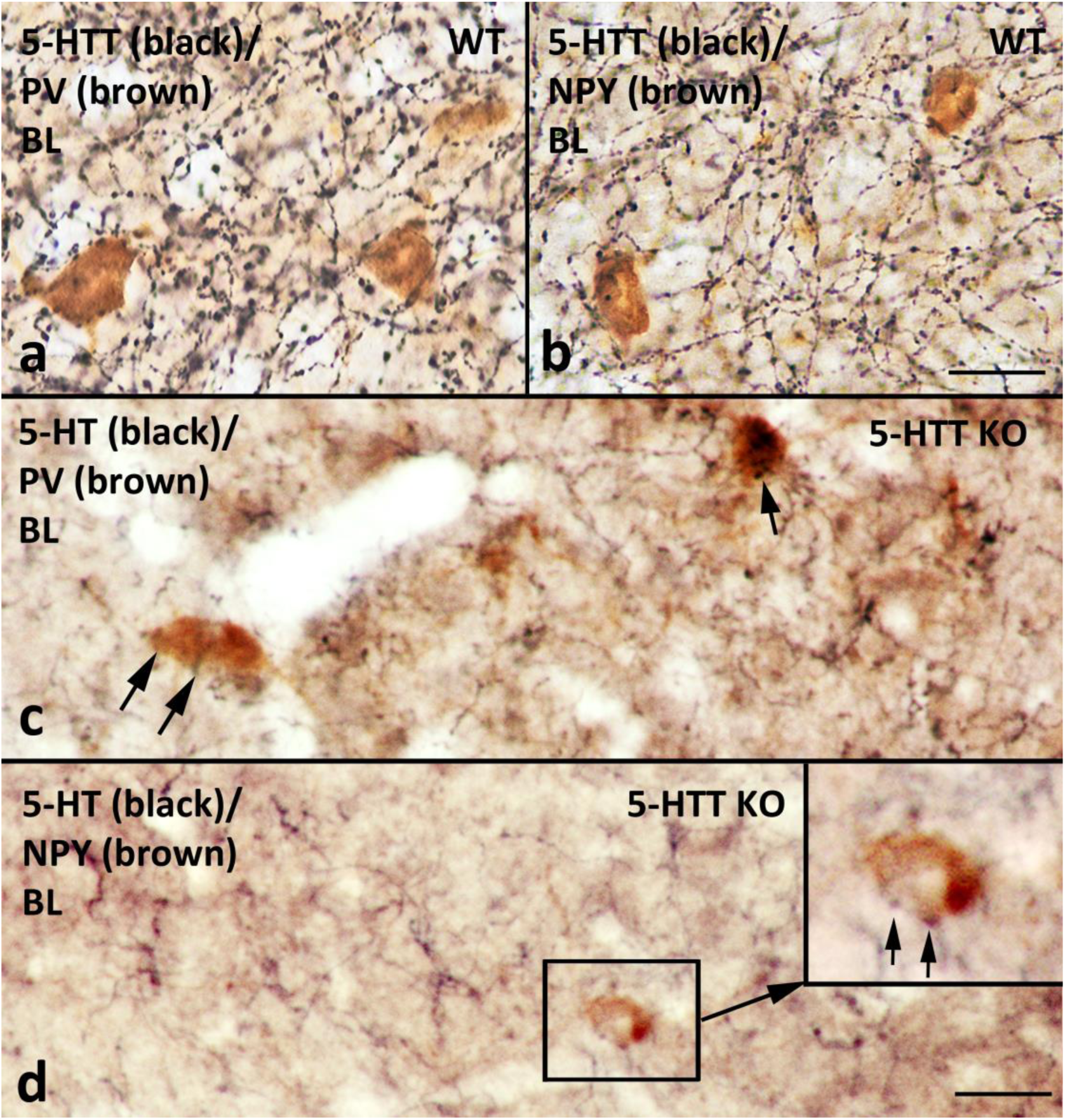
Dual immunoreactions showing perisomatic appositions of serotonergic fibers labeled by 5-HTT antibodies in the WT **(a, b)** and by 5-HT antibodies in 5-HTT KO mice **(c, d)** on PV **(a,c)** and NPY-ir neurons **(b,d)** in the BL. Numerous appositions are noted in all labelings (arrows). Bars in b for a, b and d for c, d: 20 µm

#### Perisomatic appositions of 5-HTT- and 5-HT-ir afferents on PV- and NPY-ir BLA neurons

Numerous perisomatic appositions on PV- and NPY-ir neurons are observed in 5-HTT/PV and 5-HTT/NPY immunoreactions (Fig. 4b, Fig. 5 a,b) as well as in 5-HT/PV and 5-HT/NPY immunoreactions in 5-HTT-KO mice (Fig. 5 c,d). In 5-HTT KO mice, PV- and NPY-ir neurons display perisomatic appositions in both the BL and La, and the majority of neurons receive multiple appositions (Fig. 5c-d). However, the interindividual variability of the 5-HT immunoreaction quality precludes quantitative analyses of 5-HT/PV and 5-HT/NPY dual labeled material. Such analyses were therefore only carried out for appositions of 5-HTT-ir fibers in the WT mice.

#### BLA innervation by TH-ir fibers

Catecholaminergic fibers in the BLA were detected by immunoreactions for TH, the rate-limiting enzyme of catecholamine biosynthesis. Highest density of TH-ir fibers are found in the BLp, less dense plexus in the BLa, moderately dense plexus in the La. The terminal fibers are thin and smooth with narrow and irregularly arranged varicosities. Distribution, density and morphology of TH-ir afferents in the 5-HTT KO mice closely resembles that found in WT (Fig. 6 a,b). Both PV- and NPY-ir neurons receive multiple appositions of TH-ir afferents in WT and 5-HTT KO mice (Fig. 6c-f; Table 1).

**Fig.6.**
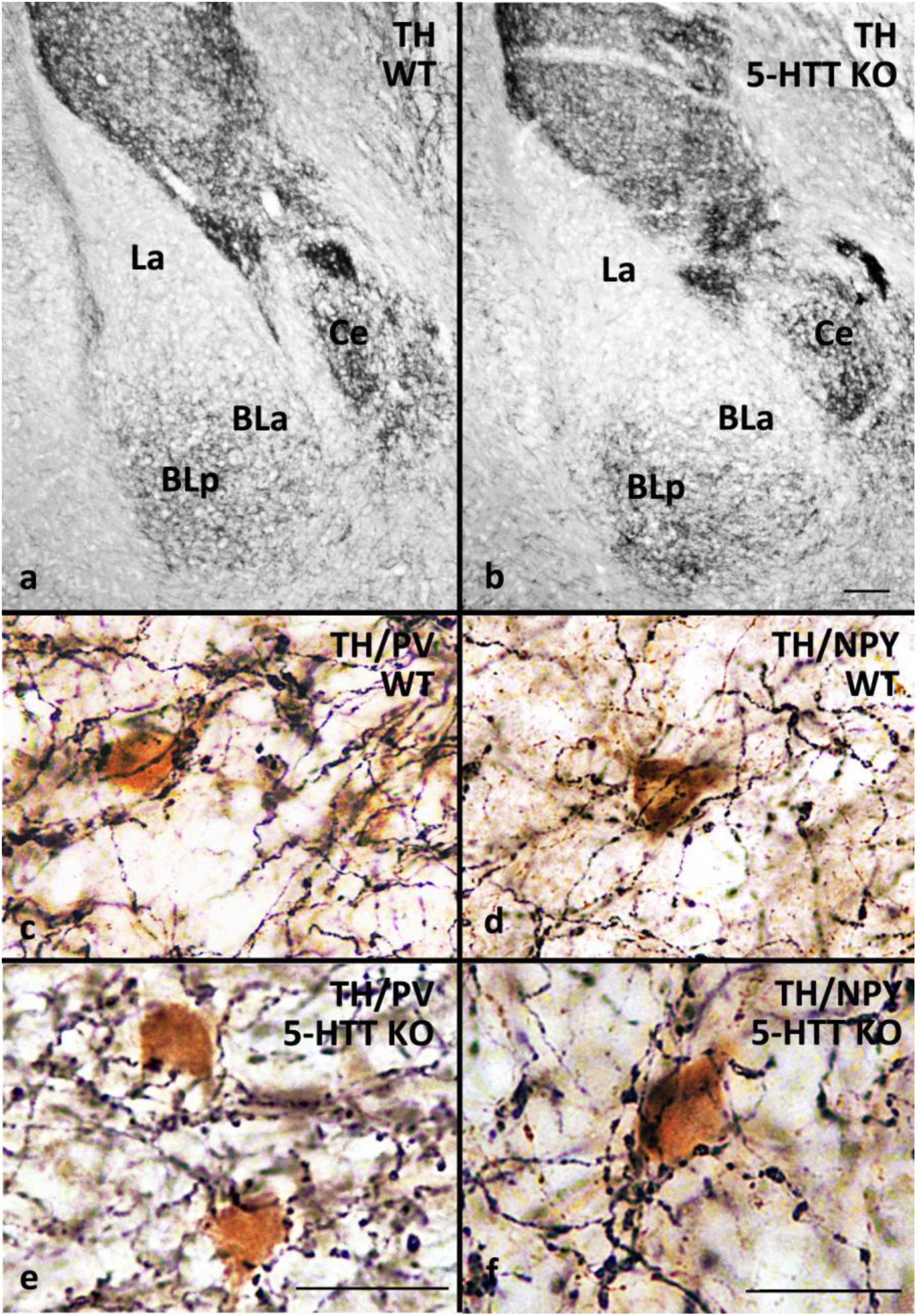
TH-ir fibers are heterogeneously distributed in the BLA of both WT **(a)** and 5-HTT KO mice **(b)**, with highest fiber density in the posterior BL (BLp), less dense plexus in the anterior BL (BLa), and scarce plexus in the La. Narrow TH-ir fibers form numerous appositions on PV-ir **(c,e)** and NPY-ir neurons **(d,f)** in both WT **(c, d)** and 5-HTT KO mice **(e,f).** Bar in b for a, b: 100 µm; Bars in e, f for c-f 25 µm.

**Table 1:**
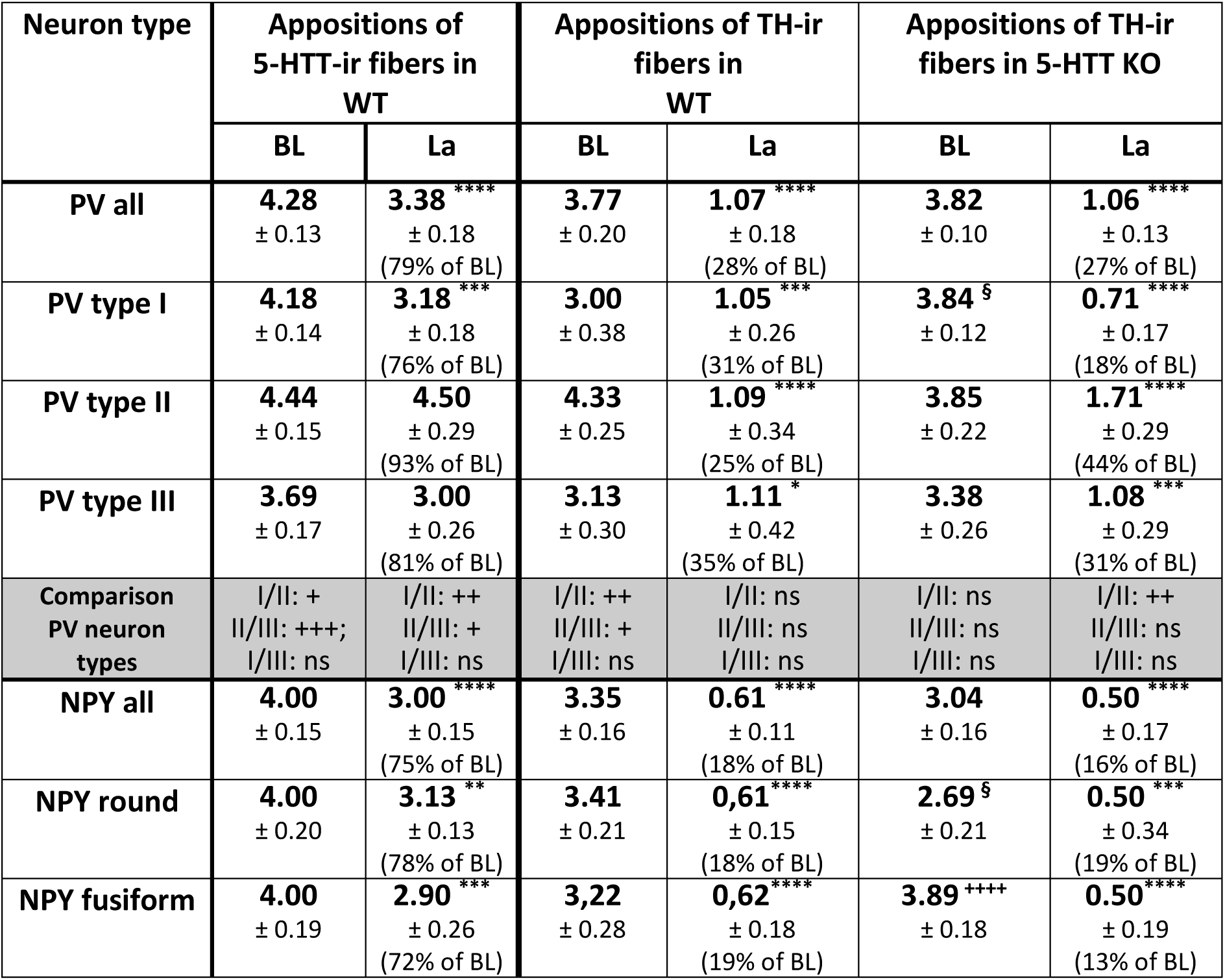
Appositions of 5-HTT-ir and of TH-ir afferents on PV- and NPY-ir neurons in the BL and La of WT and 5-HTT KO mice. Data represent mean ± SEM. Additionally, the percentages of apposition frequencies in the La compared to the BL are given for the different neuron types and afferents. Comparison BL/La: *: p<0.05; **: p<0.01; ***: p<0.001; ****: p<0.0001; Comparison of appositions on PV or NPY cell types: ^+^: p<0.05; ^++^: p<0.01; ^+++^: p<0.001; Comparison between genotypes: ^§^: p<0.05

### Quantitative assessment of perisomatic appositions of 5-HTT- and TH-ir afferents on PV and NPY-ir BLA neurons

#### Appositions of 5-HTT-ir fibers on WT PV-ir neurons (*Table 1*)

On average, PV-neurons in the BLA receive at least three perisomatic appositions by serotonergic afferents. Appositions on PV-ir neurons are significantly more numerous in the BL than in the La, with apposition frequency in the La amounting to about 80% of that observed in the BL (range 76-93% for different PV-ir neuron types). Type II PV-ir neurons receive significantly more perisomatic appositions in both nuclei than type I and III neurons.

#### Appositions of 5-HTT-ir fibers on WT NPY-ir neurons (*Table 1*)

NPY-ir neurons in the BL and La receive, on average, four and three perisomatic appositions, respectively, by 5-HTT-ir afferents. Appositions in the BL are significantly more frequent than in the La (72-78% for different NPY-ir neuron types). No statistically significant difference in apposition counts is found between round and fusiform somata. *Appositions of TH-ir fibers on PV-ir neurons (Table 1).* In both genotypes, PV-ir neurons in the BL have the highest number of perisomatic appositions. Each analyzed neuron possesses more than three appositions. In the La, perisomatic appositions on PV-ir neurons are significantly less frequent than in the BL: only 58% and 69% of the randomly selected neurons possess perisomatic appositions by TH-ir fibers in the WT and 5-HTT KO, respectively (data not shown). Apposition frequency in the La amounts to 28 and 27% of that in the BL for PV-ir neurons in the WT and the 5-HTT-KO (ranges 25-35% and 18-44%, respectively; Table 1). Type II PV-ir neurons possess significantly more appositions than type I and type III neurons in the WT BL and more appositions than type I neurons in the 5-HTT KO BL. For other nuclei and neuron types, differences in apposition frequencies do not reach statistical significance.

#### Appositions of TH-ir afferents on NPY-ir neurons (*Table 1*)

In both WT and 5-HTT KO BL, each randomly selected NPY-ir neuron displays perisomatic appositions. In the La, only 47.5% and 39%, respectively, of analyzed NPY-ir neurons show perisomatic appositions by TH-ir fibers in the WT and 5-HTT KO animals, respectively. Apposition frequency in the La amounts to only about 18% (range:18 −19%) of that in the BL in WT and 16% (range 13 – 19%) in 5-HTT KO La. Differences in apposition frequencies between the morphological subtypes (round vs. fusiform) are not observed in WT. In 5-HTT KO BL, round NPY-ir neurons possess less appositions than fusiform neurons, and less than their counterpart in the WT.

### PV- and NPY-ir neuron number and density in the BLA

Determination of the total BLA volumes analyzed in six sections across the entire rostrocaudal extent of the amygdala yielded a mean of 91.55 ± 6.67 x 10^-3^ mm³ in eight to nine WT animals and 101.2 ± 3.63 x 10^-3^ mm³ in eight 5-HTT KO mice, significant differences between the genotypes were absent (p=0.08). The same was true when volumes analyzed for the nuclei and subnuclei were compared (BL WT/KO: 60.98 ± 3.77/62.68 ± 1.6, p = 0.38; La WT/KO: 30.56 ± 3.67/37.97 ± 2.54, p = 0.12; BLa WT/KO: 48.48 ± 2.22/50.73 ± 1.69, p = 0.52; BLp WT/KO: 10.32 ± 2.04/12.49 ± 1.16, p = 0.44)

#### PV-ir neurons

Mean absolute numbers of 235.0 ± 21.8 and 220.8 ± 24.1 PV-ir neurons were recorded in the BLA for WT and 5-HTT KO mice, respectively. Neuron counts are more than 4-times higher in the BL than in the La in both genotypes, without significant genotype differences (Online resource 2). Accordingly, in the WT, the BL shows the highest density of PV-ir neurons (Fig. 7a, Online resource 2). The difference between BL and La is highly significant (p < 0.001). Within the BL, PV-ir neurons are not homogeneously distributed: BLp has a significantly higher density than BLa (Fig. 7b, Online resource 2). PV-ir neuron density increases in a rostro-caudal direction both in the BL and in the La, a significantly higher neuron density is observed at the intermediate level compared to the rostral level (Online resource 3). In 5-HTT KO animals, a similar pattern is found: PV-ir neurons are heterogeneously distributed, the differences between the nuclei and the subnuclei are significant (BL vs La; p <0.01; BLp vs BLa; p < 0.01; Fig. 7, Online resource 2). As in the WT, PV-ir neuron density increases in a rostro-caudal direction and, in the BLA and BL, is again highest at the intermediate level (Online resource 3). Although mean values of PV-ir neuron densities throughout nuclei and subnuclei in 5-HTT KO are slightly lower than in WT mice, the differences do not reach significance (Fig. 7).

**Fig. 7:**
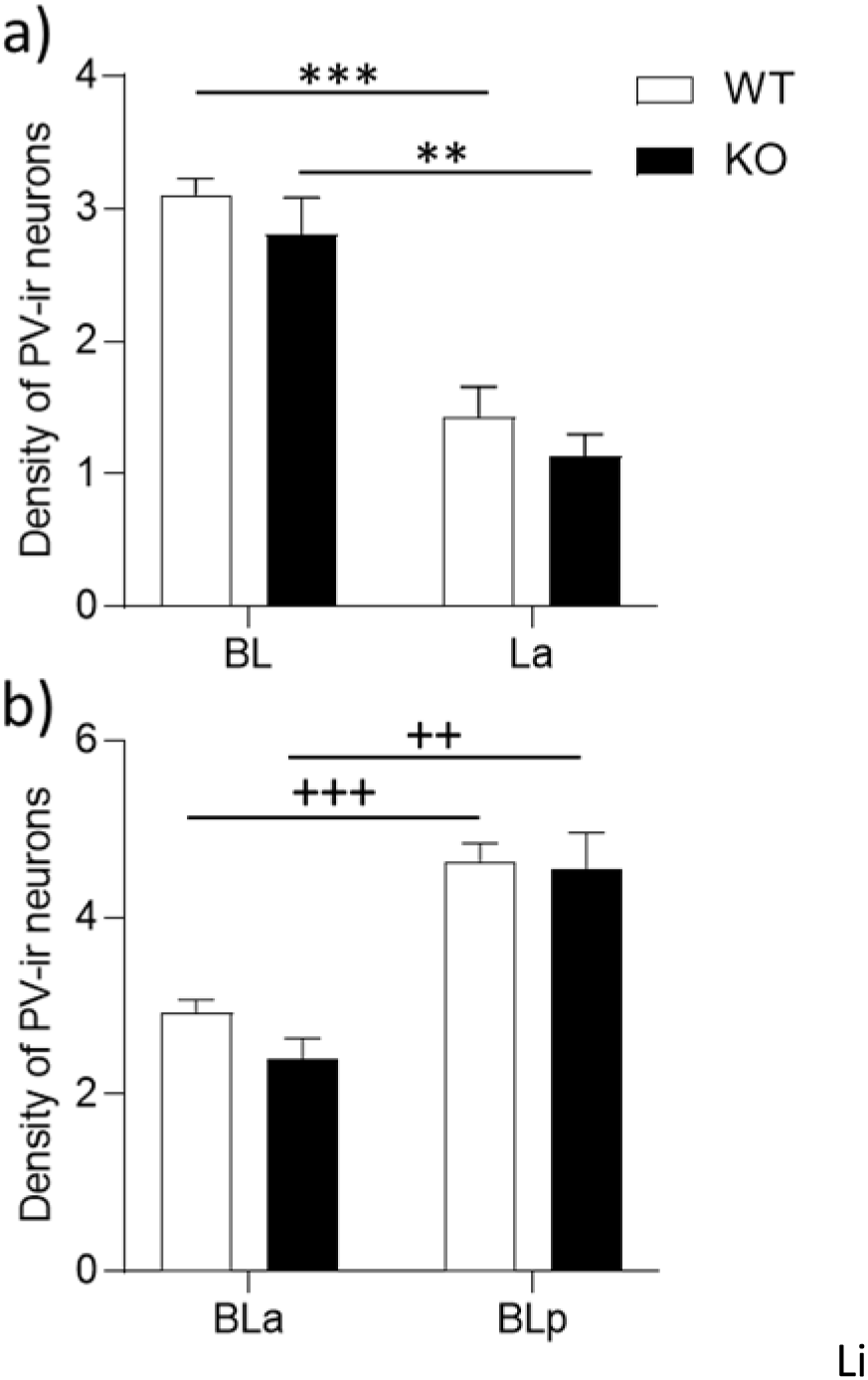
**(a)** PV-ir neuron density (number x 0.001 mm^-3^) is higher in the BL than in the La, and **(b)** is higher in the BLp than in the BLa in both genotypes. There are no significant genotype-dependent differences in these nuclei and subnuclei.

#### NPY-ir neurons

A mean total count of 127.4 ± 7.1 and 75.63 ± 5.42 NPY-ir neurons was recorded in the BLA of WT and 5-HTT KO amygdalar sections, respectively. The genotype difference in absolute numbers is highly significant (Online resource 4; p < 0.001). Neuron counts are higher in both genotypes in the BL than in the La, and for both nuclei the genotype difference is highly significant. NPY-ir neurons were additionally classified into the two morphological subtypes (Online resource 4). There are slightly more round than fusiform neurons in all nuclei in both genotypes and there are significantly less NPY-neurons of both types found in KO mice.

In the WT, the density of NPY-ir neurons is higher in the BL than in the La (Online resource 4), but the difference does not reach statistical significance. Comparing different rostro-caudal levels of the amygdala, it is noted that the density of NPY-ir neurons is highest in the rostral segment and decreases in a caudal direction (Fig. 8b; Online resource 5). This is true for both neuronal subtypes (Online resource 5). In 5-HTT-KO mice, the density seems to be also slightly higher for all morphological subtypes in the BL than in the La (Online resource 4), here again the difference within the genotype does not reach statistical significance. As in the WT, NPY-ir neuron density in the 5-HTT KO mice is highest in rostral amygdalar regions. A significantly decreased density is observed for NPY-ir neurons in all nuclei and regions of the BLA in 5-HTT KO mice compared to WT (Fig. 8, Online resource 5). Additionally, the proportion of round NPY-ir neurons compared to fusiform ones is increased in BL and La of 5-HTT KO compared to WT mice. This increase is significant when comparing the entire BLA (WT: 55.8% round, 5-HTT KO: 62.81% round; p < 0.05).

**Fig. 8:**
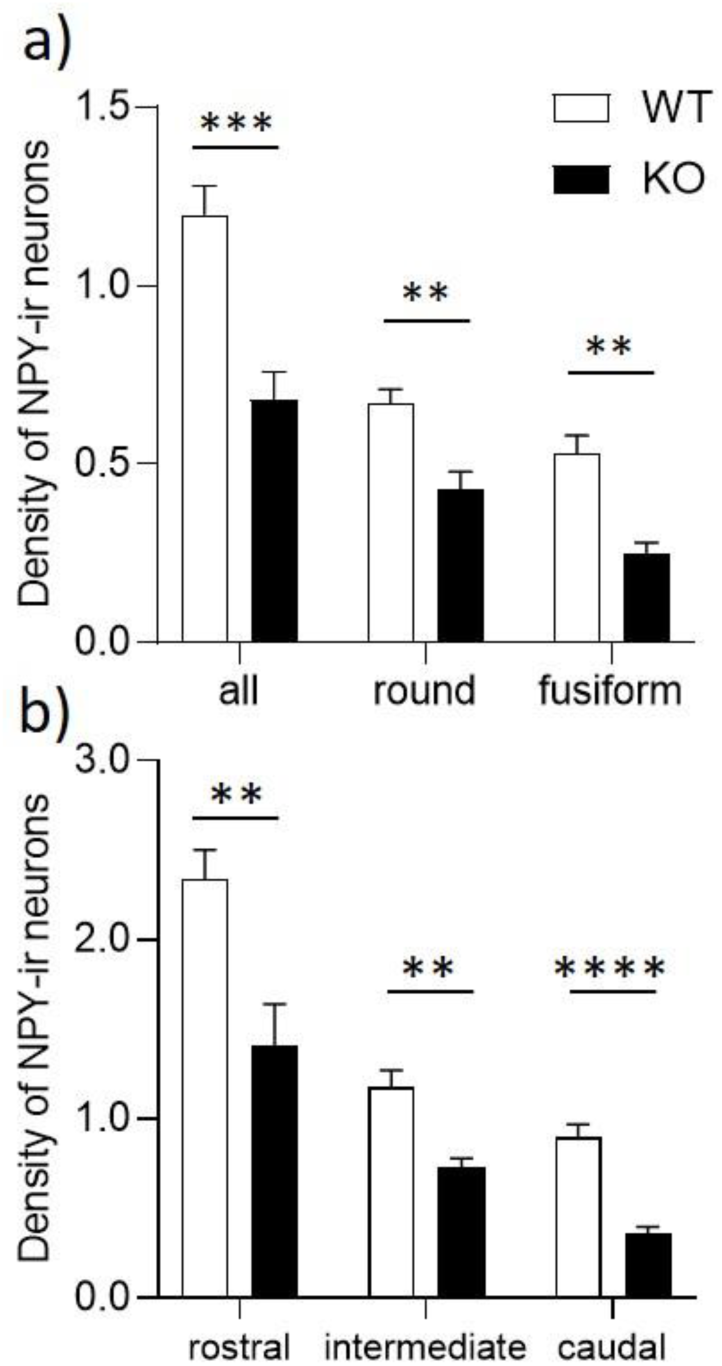
**(a)** *5-Htt* genotype-dependent density differences are observed in the BLA for all NPY-ir neurons and for round and fusiform subtypes. NPY-ir neuron density decreases within the amygdala in a rostro-caudal direction **(b)**. Genotype differences are highly significant in all subregions.

### Gene expression studies

To assess possible differences in expression levels of *Npy*, *Som*, and of the anxiogenic peptide *Crf* and their receptors (*Npyr1* and *-2*, *Sstr2* and *-4*, *Crfr1* and *-2*), qPCR analyses were carried out on laser-microdissected BLA tissue from mice of the different genotypes. Additionally, expression levels of 5-HT receptors *5Htr1a*, *-2a* and -*2c* were studied. Significant increases were found for *Npyr2*, *Sstr4* and *Crf1*. The mRNA coding for the three peptides showed no differences. Changes were also not observed in the expression levels of the three investigated 5-HT receptors (Fig. 9; Online resource 6).

**Fig. 9:**
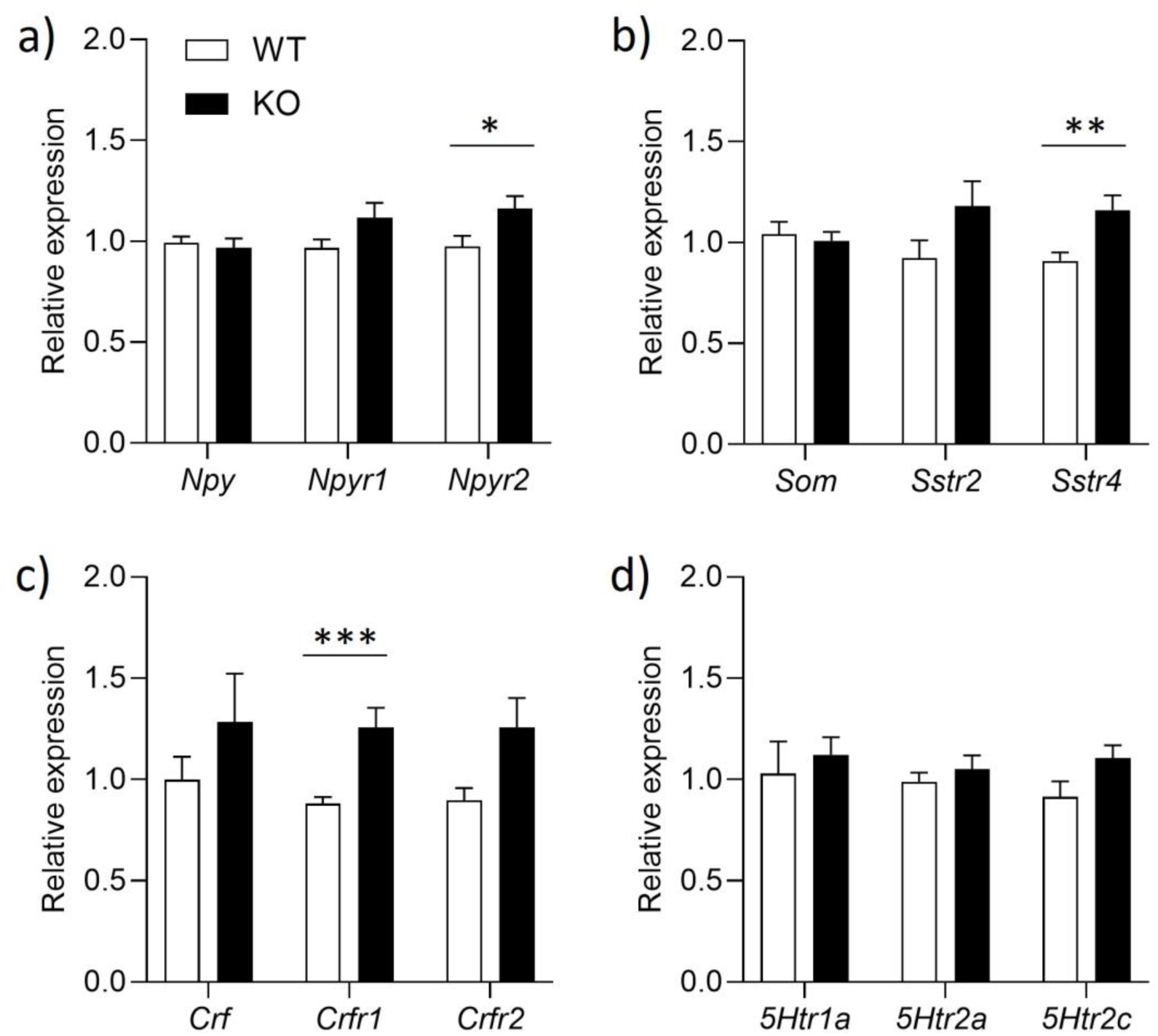
Expression levels of genes related to the NPY, SOM, CRF and 5-HT system in the BLA of WT and 5-HTT KO mice. Expression was assessed by qPCR in microdissected BLA tissue. Significant increases in relative expression levels of *Npyr2* **(a)**, *Sstr4* **(b)**, *Crfr1* **(c)** are detected in 5-HTT KO compared to WT mice. Relative expression levels of the mRNA coding for the peptides NPY, SOM, and CRF did not show alterations. Relative expression levels of serotonin receptors *5Htr1a*, -*2a* and *-2c* were also unchanged **(d).**

## Discussion

The present study documents that life-long 5-HT imbalance affects non-pyramidal neurons of the BLA: compared to WT mice, the density of neurons in which the “anxiolytic” peptide NPY (Kask et al. 2002; Truitt et al. 2009; Yilmazer-Hanke et al. 2004) is detectable by immunolabeling is significantly reduced in 5-HTT KO mice, a mouse model for anxiety- and stress-related disorders. In contrast, the density of PV-ir neurons appears unaltered. These findings add to the growing body of evidence implicating inhibitory networks of the BLA, particularly those involving the NPY-producing NPNs of the BL and La, as important players in the modulation of emotional processing and anxiety (Yilmazer-Hanke et al. 2002; Truitt et al. 2009; Bonn et al. 2013; Prager et al. 2016; Verezcki et al. 2021).

The results additionally show similarities between WT mice and published data from rats with respect to both the morphology and distribution patterns of BLA PV- and NPY-ir neurons, and with respect to the general distribution patterns of serotonergic afferents to the BLA (Kemppainen et al. 2000, McDonald 1989, Asan et al. 2013). The same holds true for the distribution patterns of fibers detected by TH immunoreactions (Asan 1997). TH is the rate limiting enzyme of catecholamine biosynthesis and thus can be found in all catecholaminergic structures. However, in the BLA of rats, TH immunoreactions have been shown to preferentially label dopaminergic fibers (Asan 1993), and TH is taken to be a dopaminergic marker also in the mouse BLA (Kalinowsky et al. 2023). The innervation patterns for serotonergic and TH-ir, presumably mostly dopaminergic, afferents appear largely conserved in 5-HTT-deficient mice. Also, in both WT and 5-HTT-deficient mice, BLA PV- and NPY-ir NPNs are perisomatically contacted by serotonergic and by TH-ir afferents. With the exception of appositions of TH-ir fibers on small round NPY neurons, no significant differences are found between the genotypes concerning the frequency of dopaminergic perisomatic contacts on neuronal subtypes. On the gene expression level, significantly increased mRNA levels are observed in the 5-HTT KO mice for specific peptide receptors (*Npyr2*, *Sstr4* and *Crfr1*), while levels of mRNA coding for the peptides, for other peptidergic receptors (*Npyr1*, *Sstr2* and *Crfr2*) and for serotonin receptors (*5Htr1a*, *-2a* and *-2c*) do not show differences between the genotypes.

### PV- and NPY-ir neurons in the WT: morphology, colocalization of PV and peptides and interrelations with monoaminergic afferents

#### Morphology and colocalization

The morphological characteristics of PV-ir BLA neurons documented in WT mice in the present study are similar to what has been previously described in rats (Kemppainen et al. 2000). There was no colocalization observed of PV- and NPY-IR. In SOM/NPY dual labelings, numerous double- and single-labeled cells were found. Immunoenzyme labeling for NPY was often restricted to “cytoplasmic streets”, presumably representing cisterns of the Golgi apparatus (Asan 1997; Takagi et al. 1983). Dual immunofluorescence showed that SOM- and NPY-immunolabeling largely overlapped in cytoplasmic compartments in double-labeled cells. SOM/NPY- and SOM-ir NPNs displayed mostly oval somata, single NPY-ir neuronal perikarya were mostly small and round, and thus morphologically resemble neurogliaform cells (Manko et al. 2012; Overstreet-Wadiche and McBain 2015; Ozsvar et al. 2024). These neurons possess small spherical cell bodies, and have been reported to usually lack SOM IR (Perrenoud et al. 2012; Vereszki et al. 2021). Vereszki et al. (2021) found that almost half (ca. 45%) of NPY reporter transgene expressing neurons were neurogliaform cells in the BLA, a percentage similar to that found for small round NPY-ir neurons in our immunoenzyme labeled material (55%) in the WT. Thus, although it has to be cautioned that specific identification of neurogliaform cells was not carried out in our investigation, it appears reasonable to assume that many of the small, round NPY-ir cells in our material are neurogliaform cells. In the rat BL, almost complete colocalization of SOM in NPY-ir BLA neurons was reported (McDonald 1989), indicating that in this species neurogliaform cells might either not be as prevalent in the BLA as in mice, or produce less NPY. All morphological characteristics found in the WT mouse in our material were preserved in 5-HTT KO mice.

#### Interrelations with 5-HTT- and TH-ir afferents

In the rat, PV-ir neurons have been shown to receive numerous perisomatic synapses, many of which are symmetric (Müller et al. 2005). Electron microscopy showed that narrow varicosities of serotonergic and dopaminergic afferents form multiple appositions on PV-ir perikarya, and small synaptic contacts were identified in most of them (Müller et al. 2007b; Pinard et al. 2008). The same holds true for NPY-ir neurons in the rat BLA, which are extensively contacted by serotonergic and TH-ir afferents at their perikarya, and small synaptic contacts have been detected (Asan et al. 1998; Bonn et al. 2013). Our findings document the presence of multiple perisomatic appositions of both monoaminergic afferent types on all morphological subgroups of PV-ir and NPY-ir BLA neurons also in the mouse. Monoamines are known to mediate their effects not only via synaptic, but also via volume transmission (Bunin and Wightman 1998; Zoli et al. 1999; Fuxe et al. 2007; Özcete et al. 2024). Although monoaminergic afferents also form contacts with dendrites and/or axon terminals (Asan 1998; Müller et al. 2007b; Bonn et al. 2013), their perisomatic appositions are in a particularly powerful position to influence the activity of their target neurons either directly or via binding to presynaptic receptors on terminals of other (mostly GABAergic) neurons forming contacts on the same somata.

Our quantitative data indicate that apposition frequencies of *serotonergic afferents* on PV- and NPY-ir somata are correlated with the 5-HTT-ir fiber densities observed in the WT BL and La, respectively, and with the soma size of the contacted neurons. The reduction in apposition frequency of serotonergic afferents on PV-ir neurons in the La as compared to the BL (to ca. 79%, Table 1) agrees with differences observed qualitatively for the serotonergic fiber densities in the two nuclei. None of the PV-ir neuron subtypes seems to be a preferential target of serotonergic afferents. It appears rather that appositions are relatively homogeneously distributed on the available soma surface of the respective target neurons. Moreover, both perisomatic apposition frequencies, and the reduction in apposition frequencies in the La compared to the BL, are similar for PV- and NPY-ir neurons (the latter to ca. 75%) in the WT, suggesting that serotonergic afferents do not prefer one of these NPN types as targets over the other in the La. Unfortunately, due to the inconsistent results for 5-HT stainings in our material, it was not possible to quantitatively assess serotonergic perisomatic appositions on PV- and NPY-ir neurons also in the 5-HTT KO mice. However, in well-stained sections, the 5-HT-ir fiber density was comparable to that of 5-HTT-immunoreactions in WT mice, and numerous perisomatic appositions were observed on both types of neurons studied also in the 5-HTT KO mice, indicating that interrelations between the serotonergic afferents and their targets are similar in the BLA of the two *5-Htt* genotypes.

PV- and NPY-ir neurons are also densely contacted by *TH-ir, presumably predominantly dopaminergic afferents* in the BL. The number of perisomatic appositions in the La amounts to about 30% of that found in the BL for PV-ir neurons, and to less than 20% for NPY-ir neurons, indicating that the comparatively scarce TH-ir innervation of the La is directed more towards PV-ir than NPY-ir neurons. Previous ultrastructural studies have documented that PV-ir neurons, but not calretinin-immunoreactive neurons, were prominent targets of dopaminergic terminals in the basolateral complex of the rat (Pinard et al. 2008). Thus, at least among NPNs of the La, PV-ir neurons appear to be a preferential target of dopaminergic afferents in both species. In this context it is interesting to note that dopamine has been shown to suppress feed-forward inhibition in the BLA (Bissière et al. 2003), a function which could well be mediated by dopaminergic innervation of PV-ir neurons. The findings described for the innervation of PV- and NPY-neurons by TH-ir fibers in the WT mouse BLA also apply to the 5-HTT KO mice. Genotype differences are, however, apparent in the innervation of NPY-ir neuronal subtypes. Thus, while appositions of TH-ir fibers are about equally frequent on the two NPY-ir neuron subtypes (round and fusiform) in the WT BL, appositions on round neurons are less frequent on round than on fusiform neurons in the 5-HTT KO, and are also less frequent than on their counterparts in the WT BL (Table 1).

### Quantitative analyses of PV- and NPY-ir neuron numbers in the BLA in WT and 5-HTT KO mice

The quantitative analyses performed in the present study yielded a number of significant findings. First, evaluation of the nuclear and subnuclear volumes analyzed in six sections throughout the rostro-caudal extent of the amygdala in eight to nine adult mice of each group yielded no genotype-dependent difference, indicating that 5-HTT deficiency did not affect the size of the BLA.

In absolute counts in WT mice, there were 1.85 times more PV-ir neurons than NPY-ir neurons recorded in the entire BLA. In the BL, PV-ir neurons counts were 2 times higher than NPY-ir neuron counts, in the La 1.3 times. In rats, PV-ir neurons account for about 40% of GABAergic non-pyramidal neurons in the BL, and for 19% in the La (McDonald and Mascagni, 2002). The percentage of NPY neurons among GABAergic neurons has not been reported in rats, but a higher percentage was found in the BL than in the La in mice by Vereszki et al. (2021). Moreover, using a combination of transgenic and viral strategies to label different types of GABAergic neurons in the mouse BLA, Vereszki and coworkers (2021) reported that 24% of NPNs expressing an NPY-reporter protein coexpress PV, 30% coexpress SOM, and 45% express neither of these markers and are considered to be neurogliaform cells. These findings differ with ours regarding the coexpression of PV and NPY, which was never found in our material. It may be that immunolabeling is too insensitive to enable detection of low amounts of peptide. This problem is frequently encountered when studying mice. For instance, CRF-producing neurons in the central amygdaloid nucleus (Ce) cannot be visualized in mice by peptide immunolabeling, although neurons detected by Crf mRNA *in situ* hybridization are as numerous in the mouse Ce as in the rat Ce (Asan et al. 2005). However, the Ce CRF-neurons are projection neurons, which rapidly transport the peptide into their long axonal processes and thus in mice obviously do not display high perikaryal peptide content, while peptidergic INs, which usually show a somatodendritic localization of the peptides in both species, can often be detected by immunolabeling in mice as in rats. Still, the immunoenzyme and immunofluorescence labeling approaches to assess the NPY content of NPNs we chose in the present study are presumably less sensitive than detection of reporter proteins in neurons capable of transcribing and translating the different peptides or proteins in transgenic mice. However, the relatively direct immunodetection of the antigens circumvents questions of expression specificity, and enables detection of the intracellular content of the peptide or protein in question under relatively physiological conditions. In any case, the number and neurochemical identity of NPY-ir neurons detected in the present study in both genotypes may represent only neurons with a rather high content of the peptide. Therefore, the term NPY-ir should be taken to represent neurons with perikaryal NPY-levels detectable by immunolabeling.

#### Density of PV-ir and NPY-ir neurons

The highest density of *PV-ir neurons* was found in the BLp, followed by the BLa and the La. PV-ir neuron density in the entire BLA increased in a rostrocaudal direction in all subnuclei, with highest density of neurons at the intermediate rostrocaudal level (−1.5 to −2.2 mm from bregma). This agrees well with the observation of Vereszki et al. (2021), who reported that the proportion of inhibitory cells among all neurons of the BLA peaked at −2.1 mm from bregma. There were no significant differences observed between the genotypes, indicating that 5-HTT deficiency does not influence PV-neuron density and distribution.

In WT mice, *NPY-ir neuron* density was slightly higher in the BL than in the La and decreased in a rostro-caudal direction. As was observed for the PV-ir neurons analyzed in the same amygdalae, the relative proportions concerning NPY-ir neuron densities between nuclei and regions were preserved in 5-HTT KO animals. However, in contrast to PV-ir neurons, there was a significant reduction found for neurons with detectable NPY-IR observed in the 5-HTT KO animals throughout the BLA. In the WT, round and fusiform NPY-ir neurons accounted for about the same proportions in BL and La, with around 55% round neurons in both nuclei. This relationship was changed in 5-HTT KO mice, with a slightly but significantly higher proportion of round neurons in the entire complex. Together with the fact that round NPY-ir neurons in the BL received significantly less perisomatic appositions of TH-ir fibers in the 5-HTT KO BL than in the WT BL, and following the arguments on possible identity of NPY-ir neurons stated above, this indicates that life-long serotonin homeostasis imbalance may differentially affect NPY production and perikaryal appositions by TH-ir fibers in different NPY-IN types of the 5-HTT KO mice. The reduction in numbers of neurons with detectable perikaryal NPY content could either be caused by a reduction in neurons capable of producing NPY in 5-HTT deficiency, possibly caused by a defect in development or developmental migration of these neurons, or by a reduction of NPY production in the different cell types.

### Expression of anxiety-related peptides and receptors

Gene expression studies revealed that there was neither a change for *Npy* nor for *Som* or *Crf* mRNA in 5-HTT KO animals. This argues against the proposition that the reduction in the number of BLA NPNs with immunohistochemically detectable NPY-IR in the 5-HTT mice is due to a dysfunctional development and/or developmental migration of NPY-NPNs. It appears rather that, in 5-HTT-deficient animals, the translation of the peptide is reduced in BLA-NPNs which transcribe its mRNA and are, in principle, capable of producing NPY, which they do in the WT. Interestingly, a recent study documented a reduction of NPY-ir NPN numbers in the BLA of aged rats. This could be reversed by intracerebroventricular injection of NGF, which led to an increased density of cholinergic varicosities in the BLA, indicating that the production of NPY peptide in BLA NPNs is regulated by extrinsic, presumably cholinergic afferents (Pereira et al. 2024). The observation that NPY appears to be localized in Golgi-like compartments in some neurons in our material suggests that the respective neurons are engaged in production of the peptide. It remains to be analyzed whether NPNs in 5-HTT KO mice can increase their NPY production on demand, perhaps in stressful situations, and, if so, how this demand is relayed to the neurons. As described above, NPY production may be regulated by extrinsic innervation from cholinergic systems (Pereira et al. 2024); the differences in apposition frequency of TH-ir fibers on NPY-ir neurons observed in the present study implicate also catecholaminergic systems (as inducers or inhibitors of NPY-production). In rats, NPY-producing neurons express, in mostly separate subpopulations, inhibitory 5-HT1A and excitatory 5-HT2C receptors (Bonn et al. 2013). 5-HT receptor expression in the different NPY-ir neurons of the mouse BLA has not been studied yet, but if there are similarities in receptor expression as there are in other features of NPY-producing neurons between rats and mice, serotonin imbalance could well exert distinct effects on BLA-NPY-neurons. In any case, increased mRNA levels of *Npyr2* and *Crfr1*, receptors that mediate anxiogenic effects (Giesbrecht et al. 2010), and of *Sstr4*, which presumably mediates anxiolysis (Scheich et al. 2016), support the observation that the BLA-NPY-system is indeed affected by life-long 5-HT imbalance, and additionally show that the CRF- and SOM-systems are also changed in 5-HTT KO mice.

Of note, the absence of alterations in PV-ir neuron numbers in 5-HTT KO mice does not exclude effects on PV-ir neuron functions. Indeed, overexpression of 5-HTT in mice has been documented to result in reduced recruitment of PV-ir neurons in BL and La and in impaired fear-related behavior (Bocchio et al. 2015). PV neurons express the 5-HT2A receptor, and electrophysiological studies indicated that 5-HT-induced activation of PV-ir neurons and thus basic inhibition of principal cells was impaired due to reduction in 5-HT2A receptor function in 5-HTT overexpressing mice (Bocchio et al. 2015). Also, the finding that mRNA of the 5-HT receptors analyzed in the present study did not display differences in 5-HTT KO compared to WT mice does not argue against a possible dysfunctionality of receptor-mediated signal transduction, since transduction efficiency of receptors does not only depend on the amount of receptor present, but also on molecular interactions with various other membrane or cytoplasmic molecules (Bohn et al. 2010, Chruscika et al. 2019, Xu et al. 2021).This, of course, also applies to the other receptors analyzed in the present study.

### Functional implications

Analyses of the synaptology have documented that networks formed by different populations of GABAergic NPNs fulfill distinct tasks in the regulation of pyramidal cell activity and information processing in BL and La. Synapses of PV-ir neurons, located mainly perisomatically, provide an extensive inhibitory stimulus resulting in a constant, basic control of the activity of the pyramidal cells. The synaptic contacts of SOM-ir INs, and thus most likely of many NPY-ir INs coexpressing SOM, are located at distal dendrites of pyramidal cells. These INs are ideally positioned to modulate excitatory synaptic transmission also directed mainly at distal dendrites and spines of PN, and thus may influence dendritic plasticity of PNs (Krabbe et al. 2018). NPY-ir neurogliaform cells are considered one of the major contributors to slow-phasic and tonic GABA release in the BLA, resulting in both GABA-A and GABA-B receptor mediated inhibition of PNs. However, their exact contribution to BLA circuit operation remains unknown (Ozsvar et al. 2024), as are possible effects of NPY released from these neurons. Investigations of the activity and/or of the NPY-production in BLA-INs of 5-HTT KO mice under different conditions may yield insights into these questions.

Neuroplasticity allows adaptation to changed environmental conditions. The BLA, particularly the La, is considered to be the main input station of the amygdala. Various informations from cortical and subcortical structures are integrated here. Thus, it is not surprising that the BLA is a place of intense neuroplasticity. Most classic studies of conditioned fear focus on NMDA- and AMPA-receptor-dependent long-term potentiation (LTP) of glutamatergic projection neurons, but in recent years it has become increasingly obvious that GABAergic neurons play a decisive role in this process as well (Ehrlich et al. 2009; Prager et al. 2016).

In 5-HTT KO mice, number and density of spines on distal dendrites of La and BL PNs are significantly higher than in WT animals (Wellman et al. 2007; Nietzer et al. 2011). This has been suggested to be a morphological correlate for an increased excitability of the BLA-PNs and, as a consequence, the *per se* anxious phenotype of these animals. Based on the findings of the present study, it could be argued that a lack of NPY transmission from SOM/NPY-producing INs and/or from neurogliaform cells may lead to a reduced inhibition and thus an increased excitatory drive on PN dendrites, which results in an increased spine density. In addition, Nietzer et al. (2011) showed that stress in WT mice leads to a significant increase in spine density to the level observed in naïve 5-HTT KO mice, whereas no further increase was observed in stressed 5-HTT KO mice, indicating a lack of neuroplasticity. In this respect, following suggestions by Mackay et al. 2019, it could be argued that in the WT mice NPY, released from GABAergic IN terminals on distal dendrites and acting on presynaptic NPYR2 autoreceptors of these terminals, may induce selective disinhibition of dendritic domains under stress, enabling situation-adequate dendritic plasticity as a “memory trace” of stressful events. The relative scarcity of NPY in 5-HTT mice, which is reflected in an increased production of the *Npyr2* mRNA and possibly protein, could preclude such plasticity. At the same time, fear responses were significantly increased in stressed 5-HTT KO mice compared to stressed WT controls (Holmes et al. 2003b, Karabeg et al. 2013, Hohoff et al. 2013). The dysfunctional inhibitory drive in 5-HTT KO mice may be additionally compromised due to stress induced alterations, for instance in monoaminergic transmission in the BLA, resulting in an inability to adequately modulate hyperexcitation of pyramidal cells and consequently leading to the observed increased fear-related behavior. The hypotheses generated from the findings of the present work will have to be tested in future investigations, e.g. of the innervation characteristics of NPY-ir neurons of the BLA as well as the density of SOM-ir neurons in 5-HTT-deficient mice.

## Conclusion

Our findings show that the disrupted 5-HT homeostasis in 5-HTT KO mice leads to alterations of specific inhibitory networks in La and BL. Disturbances in the complex interplay of inhibition and disinhibition/excitation may be the result. Alterations in the network formed by NPY-ir neurons are clearly of particular physiological and clinical relevance. Similar changes in individuals who are carriers of the *S*-allele of the *5-HTT* gene with a resulting lower functional expression of 5-HTT could be contributing to the increased activity and responsiveness of the amygdala and, ultimately, for the altered processing of and response to emotional stimuli in these individuals.

## Acknowledgements

We are indepted to Rita Herrmann, Karin Reinfurt, Sieglinde Schenk, and Gabriela Ortega for invaluable technical assistance. The authors are grateful to Herbert Schwegler for helpful discussions. The work was supported by the DFG (SFB581, Z3 to EA and B6 to KPL, and RTG 1253 to EA, KPL and ASB, and RTG 2660 C3 to EA and ASB; SFB TRR58, A01 and A05 to KPL and ASB). KPL was also supported by the EU Horizon 2020 Research and Innovation Programme under Grant No. 728018 (Eat2beNICE), and Grant No. 953327 (Serotonin and Beyond).

Results of the present study are part of the published dissertation work by H. Schwert (Schwert et al., 2017).

## Data availability statement

All data supporting the findings of this study are available within the paper and its Supplementary Information.

