## Supplemental tables 1-6 for "Impact of serotonin transporter deficiency on parvalbumin- and neuropeptide Y-producing interneurons of the basolateral amygdala"

### Online Resource 1

Primer targets and their specifications. F: Forward primer sequence; R: Reverse primer sequence. Manufactured by Sigma Aldrich at 100 nM. bp: base pairs.

| Gene | Symbol | Product [bp] | Intron spanning | Accession number/Reference sequence | Primer sequence |
|---|---|---|---|---|---|
| Reference genes | | | | | |
| Actin beta | *Actb* | 84 | no | NM_007393.5 | F: ATGTGGATCAGCAAGCAGGA<br>R: AGCTCAGTAACAGTCCGCCTA |
| Beta 2 microglobulin | *B2m* | 126 | yes | NM_009735.3 | F: ACCGTCTACTGGGATCGAGA<br>R: TGCTATTTCTTTCTGCGTGCAT |
| Glyceraldehyde 3-phosphate dehydrogenase | *Gapdh* | 135 | no | NM_008084 | F: GTGATGGGTGTGAACCACGA<br>R: GGTCATGAGCCCTTCCACAA |
| GDP dissociation inhibitor beta | *Gdi2* | 126 | no | NM_008112 | F: GTCAGAATTGGTTGGTTCTGTTC<br>R: AGCTCTTGGATCACACAATCG |
| 60S acidic ribosomal protein P0 | *Rplp0* | 83 | yes | NM_007475 | F: GAGGCCACACTGCTGAACAT<br>R: ATGCTGCCGTTGTCAAACAC |
| Target genes | | | | | |
| Serotonin 1A receptor | *5Htr1a* | 144 | yes | NM_008308.4 | F: GATCTCGCTCACTTGGCTCA<br>R: AAAGCGCCGAAAGTGGAGTA |

| | | | | | |
|---|---|---|---|---|---|
| Serotonin 2A receptor | *5Htr2a* | 112 | yes | NM_172812.3 | F: CCATAGCCGCTTCAACTCCA<br>R: CGAATCATCCTGTAGCCCGA |
| Serotonin 2C receptor | *5Htr2c* | 73 | yes | NM_008312 | F: GCAATAATGGTGAACCTGGGC<br>R:ACTGCCAAACCAATAGGCCA |
| Corticotropin-releasing factor | *Crf* | 76 | no | NM_205769 | F: CAACCTCAGCCGGTTCTGAT<br>R: CAGCGGGACTTCTGTTGAGA |
| Corticotropin-releasing factor receptor 1 | *Crfr1* | 136 | yes | NM_007762 | F: GGAACCTCATCTCGGCTTTCA<br>R: GTTACGTGGAAGTAGTTGTAGGC |
| Corticotropin-releasing factor receptor 2 | *Crfr2* | 62 | yes | NM_001288618 | F: ATGACGAAGTTACGAGCATCCA<br>R: GCCTTCACTGCCTTCCTGTATT |
| Neuropeptide Y | *Npy* | 106 | yes | NM_023456 | F: CAGATACTACTCCGCTCTGCGACACTACAT<br>R: TTCCTTCATTAAGAGGTCTGAAATCAGTGTCT |
| Neuropeptide Y receptor 1 | *Npyr1* | 105 | yes | NM_010934 | F: ATTTCGGCCCACTCTGCTTT<br>R: ACCTGTACTTACTGTCCCGGA |
| Neuropeptide Y receptor 2 | *Npyr2* | 77 | yes | NM_001205099 | F: CATCTGAGAAGGAACGCGCA<br>R: CTACCGGGCCCATCTTCAGA |
| Somatostatin | *Som* | 112 | yes | NM_009215 | F: GACCCCAGACTCCGTCAGTTT<br>R: TCTCTGTCTGGTTGGGCTCG |
| Somatostatin receptor type 2 | *Sstr2* | 97 | no | NM_001042606 | F: GAGCAGTTGAATGGGAGCCA<br>R: AGTATGGCTCGGTCTGGTTG |
| Somatostatin receptor type 4 | *Sstr4* | 92 | no | NM_009219 | F: AGCGGGCATGGTCACTATC<br>R: AGCGTAGGATCACGAAGATGA |

### Online Resource 2

Mean values ± SEM of PV-ir neuron numbers and densities (x 0.001 $mm^{-3}$) of in the BLA.

Comparison between BL and La: **: p <0.01, ***: p <0.001; Comparison between BLp and BLa: **+**: p<0.05, **++**: p<0.01, **+++**: p<0.001

| Genotype | 5-HTT WT | 5-HTT KO | 5-HTT WT | 5-HTT KO |
|---|---|---|---|---|
| Nucleus | Absolute numbers | Absolute numbers | Density (x 0.001$mm^{-3}$) | Density |
| BLA | 235.0 (± 21.8) | 220.8 (± 24.1) | 2.56 ± 0.14 | 2.19 ± 0.23 |
| La | 43.75 ± 7.63 | 42.88 ± 6.52 | 1.43 ± 0.23 | 1.14 ± 0.16 |
| BL | 191.4 ± 15.81*** | 177.9 ± 18.19*** | 3.11± 0.12*** | 2.81 ± 0.27** |
| BLa | 49.00 ± 11.04 | 56.63 ± 6.60 | 2.93 ± 0.14 | 2.40 ± 0.24 |
| BLp | 143.0 ± 12.67+ | 121.3 ± 12.44++ | 4.64 ± 0.20+++ | 4.56 ± 0.40++ |

### Online Resource 3

Mean values ± SEM of PV neuron densities (neuron numbers x 0,001mm$^{-3)}$ in two sections each from the rostral (-0.80 to -1.5 mm from bregma), intermediate (-1.5 to -2.2 mm) and caudal BLA (-2.2 to -2.9 mm). Differences in densities between the genotypes are not significant. In both genotypes, differences in densities between intermediate and rostral levels are significant for the entire BLA and for the BL (§), and differences between the caudal and the rostral level are significant for the BL (*). Data are represented as mean values ± SEM. *,§: p<0.05; §§: p<0.01; ***: p<0.001.

| Nucleus/ rostrocaudal level | 5-HTT WT PV neurons | 5-HTT KO PV neurons |
|---|---|---|
| BLA rostral | 1.88 ± 0.24 | 1,56 ± 0,23 |
| BLA intermediate | 2.93 ± 0.15§§ | 2,39 ± 0,22§ |
| BLA caudal | 2.56 ± 0.22 | 2,24 ± 0,18* |
| La rostral | 1.22 ± 0.32 | 1,00 ± 0,24 |
| La intermediate | 1.48 ± 0.29 | 1,31 ± 0,23 |
| La caudal | 1.57 ± 0.23 | 1,14 ± 0,15 |
| BL rostral | 2.12 ± 0.32 | 1,75 ± 0,24 |
| BL intermediate | 3,40 ±0,24§§ | 2,92 ± 0,26§§ |
| BL caudal | 3,21 ± 0,23* | 3,11 ± 0,22*** |

### Online Resource 4

Numbers and densities (x 0.001 mm$^{-3}$) of NPY-ir neurons in the BLA in different genotypes. Data are represented as mean values ± SEM. *: p<0.05; **: p<0.01; ***: p<0.001.

| Genotype | 5-HTT WT | 5-HTT KO | | 5-HTT WT | 5-HTT KO | |
|---|---|---|---|---|---|---|
| Nucleus/ Neuron types | Absolute numbers (Subtype Percentage of all) | Absolute Numbers (Subtype Percentage of all) | p-values (genotype; absolute numbers differences) | Density | Density | p-values (genotype; density differences) |
| BLA all | 127.4 ± 7.1 | 75.63 ± 5.42 | 0.0009*** | 1.20 ± 0.08 | 0.68 ± 0.08 | 0.0010*** |
| BLA round (%of BLA all) | 71.11 ± 3.57 (55.8%)* | 47.50 ± 3.20 (62.81%)*(percentage difference between genotypes p=0.05) | 0.0024** | 0.67 ± 0.04 | 0.43 ± 0.05 | 0.0055** |
| BLA fusiform (%of BLA all) | 56.33 ± 5.09 (44.20%) | 28.13 ± 2.74 (37.19%) | 0.0017** | 0.53 ± 0.05 | 0.25 ± 0.03 | 0.0016 ** |
| La all | 33.22 ± 2.41 | 21.38 ± 1.75 | 0.0059** | 0.98 ± 0.1 | 0.58 ± 0.07 | 0.0111* |

| | | | | | | |
|---|---|---|---|---|---|---|
| La round (%of La all) | 18.11 ± 1.87 (54.52%) | 14.25 ± 1.08 (66.67%) | 0.2048 | 0.53 ± 0.07 | 0.38 ± 0.04 | 0.0927 |
| La fusiform (%of La all) | 15.11 ± 1.84 (45.48%) | 7.125 ± 1.43 (33.33%) | 0.0057** | 0.44 ± 0.07 | 0.20± 0.05 | 0.0037** |
| BL all | 94.22 ± 5.71 | 54.25 ± 4.81 | 0.0013** | 1.31 ± 0.08 | 0.73 ± 0.09 | 0.0006*** |
| BL round (% of BL all) | 53.00 ± 3.83 (56.25%) | 33.25 ± 3.32 (61.29%) | 0.0039** | 0.73 ± 0.05 | 0.45 ± 0.09 | 0.0037** |
| BL fusiform (% of BL all) | 41.22 ± 3.79 (43.75%) | 21.00 ± 1.96 (38.71%) | 0.0020** | 0.58 ± 0.06 | 0.28 ± 0.03 | 0.0010*** |

### Online Resource 5

Mean values ± SEM of NPY neuron densities (neuron numbers x 0,001mm$^{-3}$) in two sections from the rostral (-0.80 to -1.5 mm from bregma), intermediate (-1.5 to -2.2 mm) and caudal BLA (-2.2 to -2.9 mm). Most differences in the density of all NPY neurons, round NPY neurons, and fusiform NPY neurons in the BLA are significant when comparing their density in the mid-rostral levels (§), the caudal-rostral levels (+), and the mid-caudal levels (&) in both WT and 5-HTT KO. Data are represented as mean values ± SEM. §: p<0.05; §§, &&: p<0.01; &&&: p<0.001; §§§§, ++++, &&&&: p<0.0001. Comparison between genotypes, **: p<0.01, ***: p<0.001, ****: p<0.0001.

| Nucleus/ rostrocaudal level | 5-HTT WT NPY neurons x 0,001mm$^{-3}$ | 5-HTT KO NPY neurons x 0,001mm$^{-3}$ |
|---|---|---|
| all neurons rostral | 2.34 ± 0.16 | 1.41 ± 0.23** |
| all neurons intermediate | 1.18 ± 0.09§§§§ | 0.73 ± 0.05**; §§ |
| all neurons caudal | 0.90 ± 0.07++++ | 0.36 ± 0.04 ****; ++++; &&&& |
| Round neurons rostral | 1.32 ± 0.13 | 0.81 ± 0.13** |
| Round neurons intermediate | 0.69 ± 0.05§§§§ | 0.47 ± 0.04**; §§ |
| Round neurons caudal | 0.48 ± 0.05++++; && | 0.24 ± 0.004***; ++++; &&&& |
| Fusiform neurons rostral | 1.02 ± 0.09 | 0.56 ± 0.11** |
| Fusiform neurons intermediate | 0.49 ±0.06§§§§ | 0.27 ± 0.02**; § |
| Fusiform neurons caudal | 0.43 ± 0.05++++ | 0.14 ± 0.02***; ++++; &&& |

### Online Resource 6

Descriptive and explorative statistics of quantitative real-time PCR data. Expression levels in the BLA of WT and 5-HTT KO mice are shown. Pairwise comparison was performed using either the parametric T-test or the non-parametric Mann-Whitney U-test. Significant p-values are indicated by bold numbers.

| Gene | Mean (± SEM) of different genotype | | Levene's test p-value | Shapiro-Wilk Test p-value | | T-test p-value | Mann Whitney U test p-value |
|---|---|---|---|---|---|---|---|
| | 5-HTT WT (n=10) | 5-HTT KO (n=8) | | 5-HTT WT | 5-HTT KO | | |
| *5Htr1a* | 1.03 (± 0.16) | 1.12 (± 0.09) | 0.354 | **0.001** | **0.025** | | 0.101 |
| *5Htr2a* | 0.99 (± 0.05) | 1.05 (± 0.07) | 0.366 | 0.909 | 0.607 | 0.444 | |
| *5Htr2c* | 0.91 (± 0.08) | 1.11 (± 0.06) | 0.251 | 0.311 | 0.764 | 0.077 | |
| *Crf* | 0.99 (± 0.11) | 1.28 (± 0.24) | **0.026** | 0.594 | 0.229 | | 0.408 |
| *Crfr1* | 0.88 (± 0.03) | 1.26 (± 0.09) | 0.077 | 0.323 | **0.049** | | **<0.001** |
| *Crfr2* | 0.90 (± 0.06) | 1.26 (± 0.15) | **0.011** | 0.687 | 0.517 | | 0.055 |
| *Npy* | 0.99 (± 0.03) | 0.97 (± 0.04) | 0.239 | 0.420 | 0.631 | 0.932 | |
| *Npyr1* | 0.97 (± 0.04) | 1.12 (± 0.07) | 0.314 | 0. 125 | 0. 943 | 0.079 | |
| *Npyr2* | 0.97 (± 0.05) | 1.16 (± 0.06) | 0.965 | 0.221 | 0.438 | **0.036** | |
| *Som* | 1.04 (± 0.06) | 1.01 (± 0.04) | 0.137 | 0.122 | 0.569 | 0.706 | |
| *Sstr2* | 0.92 (± 0.09) | 1.18 (± 0.12) | 0.571 | 0.096 | 0.506 | 0.102 | |
| *Sstr4* | 0.91 (± 0.04) | 1.16 (± 0.07) | 0.245 | 0.916 | 0.912 | **0.006** | |
